# Multi-omic phenotyping of *MAPT* V337M neurons reveals early changes in axonogenesis and tau phosphorylation

**DOI:** 10.1101/2024.06.04.597496

**Authors:** Gregory A. Mohl, Gary Dixon, Emily Marzette, Justin McKetney, Avi J. Samelson, Carlota Pereda Serras, Julianne Jin, Nabeela Ariqat, Andrea Keys, Cristian Chavira, Andrew Li, Steven C. Boggess, Danielle L. Swaney, Martin Kampmann

## Abstract

Tau aggregation is a hallmark of several neurodegenerative diseases, including Alzheimer’s disease and frontotemporal dementia. There are disease-causing variants of the tau-encoding gene, *MAPT*, and the presence of tau aggregates is highly correlated with disease progression. However, the molecular mechanisms linking pathological tau to neuronal dysfunction are not well understood. This is in part due to an incomplete understanding of the normal functions of tau in development and aging, and how the associated molecular and cellular processes change in the context of causal disease variants of tau. To address these questions in an unbiased manner, we conducted multi-omic characterization of iPSC-derived neurons harboring the *MAPT* V337M mutation or *MAPT* knockdown. RNA-seq, ATAC-seq, and phosphoproteomics revealed that both V337M mutation and tau knockdown perturbed levels of transcripts and phosphorylation of proteins related to axonogenesis or axon morphology. When we directly measured axonogenesis, we found that both *MAPT* V337M and *MAPT* knockdown caused decreased axon length. Surprisingly, we found that neurons with V337M tau had much lower tau phosphorylation than neurons with WT tau. CRISPR-based screens uncovered regulators of tau phosphorylation in neurons and found that factors involved in axonogenesis modified tau phosphorylation in both *MAPT* WT and *MAPT* V337M neurons. Intriguingly, the p38 MAPK pathway specifically modified tau phosphorylation in *MAPT* V337M neurons. We propose that V337M tau perturbs tau phosphorylation and axon morphology pathways that are relevant to the normal function of tau in development, which could contribute to previously reported cognitive changes in preclinical *MAPT* variant carriers.

## Introduction

Neurodegenerative diseases are a growing public health burden and remain very challenging to treat because we lack a complete understanding of the underlying disease mechanisms. A common theme in many neurodegenerative diseases is the aggregation of pathological proteins [1]. Tau aggregation is a hallmark of neurodegenerative diseases collectively called tauopathies, including Alzheimer’s disease and frontotemporal dementia. In Alzheimer’s disease, tau aggregation and phosphorylation changes correlate better with disease progression than amyloid beta pathology [2] despite clear genetic evidence linking amyloid beta to the disease [3]. In frontotemporal dementia, rare causal variants of tau that are fully penetrant for the disease prove a direct role for tau in disease pathogenesis [4].

Tremendous progress has been made in revealing the diverse molecular and cellular mechanisms that are disrupted by pathogenic tau. Recent work in human induced pluripotent stem cell (iPSC)-derived neurons has shown that pathogenic variants of tau sensitize neurons to different types of cellular stress and that this effect can be rescued by lowering tau levels via autophagy [5]. Other groups have shown that tau interferes with RNA splicing and stress granules homeostasis [6–9], disrupts the nuclear envelope [10–12], perturbs axonal trafficking [13, 14] or disrupts mitochondrial dynamics [15]. Acetylated tau has also been shown to disrupt chaperone mediated autophagy, rerouting tau and other clients to be degraded by other mechanisms [16]. Pathogenic tau has also been shown to perturb plasticity of the axon initial segment and cause changes to neuronal excitability [17] and has been implicated in driving excitotoxicity [9, 18–20]. Many of these data support a tau toxic gain-of-function model, and tau lowering has been successfully shown to be beneficial in cultured neurons and animal models [5, 21]. In fact, tau lowering is currently being tested in the clinic by antibodies and ASOs [22].

These focused studies have linked tau to diverse cellular processes that go awry in neurodegeneration. However, there are few unbiased and comprehensive studies that examine phenotypes on multiple intracellular levels or with respect to normal tau function, leaving many open questions about the direct effects of pathogenic tau and how diverse cellular phenotypes interact.

To characterize the earliest changes that pathogenic tau causes in human neurons and to understand mechanistically how pathogenic tau causes human disease, we used a multi-omic approach to unbiasedly determine the cellular phenotypes linked to pathogenic tau. We modeled pathogenic tau by using human iPSC-derived neurons with the *MAPT* V337M mutation, a known cause of frontotemporal dementia. We used two sets of iPSCs, one from a healthy donor (WTC11) and one from a patient with the *MAPT* V337M mutation (GIH6C1) [17, 23].

Our RNA-seq, ATAC-seq, proteomics and phosphoproteomics results all point to changes in axonogenesis due to the *MAPT* V337M mutation. Recently published mouse phosphoproteomics datasets in tau knockout mice and P301S mice strongly support the link between tau and axonogenesis factors and intriguingly suggest that these effects are due to tau loss of function [24, 25]. We have found that tau knockdown and *MAPT* V337M mutation have overlapping effects on the levels and phosphorylation of proteins relevant to axonogenesis and axon morphology, suggesting that the mutation perturbs a normal function of tau. We found that both the *MAPT* V337M mutation and *MAPT* knockdown inhibited axonogenesis in human neurons. Early differentiated *MAPT* V337M neurons have hypophosphorylated tau, which is recapitulated by artificially overexpressing V337M tau but not WT or R406W tau in neurons with endogenous tau knockdown. Unbiased CRISPR screens for regulators of tau phosphorylation uncovered axonogenesis-related regulators of tau phosphorylation and show that the p38 MAPK pathway may play a role in modifying tau phosphorylation specifically in V337M neurons. We propose that V337M tau perturbs tau phosphorylation and axon morphology pathways that are relevant to the normal function of tau in development, which could contribute to previously reported cognitive changes in preclinical *MAPT* variant carriers.

## Results

### MAPT V337M and MAPT knockdown perturb the transcription of axonogenesis-related genes

iPSCs generated from a healthy individual (WTC11, referred to as *MAPT* WT [26]) or an FTD patient with the *MAPT* V337M mutation (GIH6C1, referred to as \**MAPT* Het [23]) were edited in previous work [17, 23] with Cas9 to generate isogenic pairs either introducing or correcting the *MAPT* V337M mutation (Fig. 1A). The *MAPT* WT iPSCs were edited with Cas9 to generate a heterozygous *MAPT* V337M/WT clone (*MAPT* Het) and homozygous *MAPT* V337M/V337M clone (*MAPT* Hom). The \**MAPT* Het iPSCs were corrected with Cas9 to generate a *MAPT* WT/WT clone (\**MAPT* WT). We engineered the GIH6C1 lines to introduce a doxycycline-inducible Ngn2 for neuronal differentiation and CRISPRi machinery. We transduced the iPSCs with lentiviral sgRNAs targeting *MAPT* to knockdown tau or non-targeting control (NTC) sgRNAs for further mechanistic characterization (Figure S1A-C).

**Figure 1:**
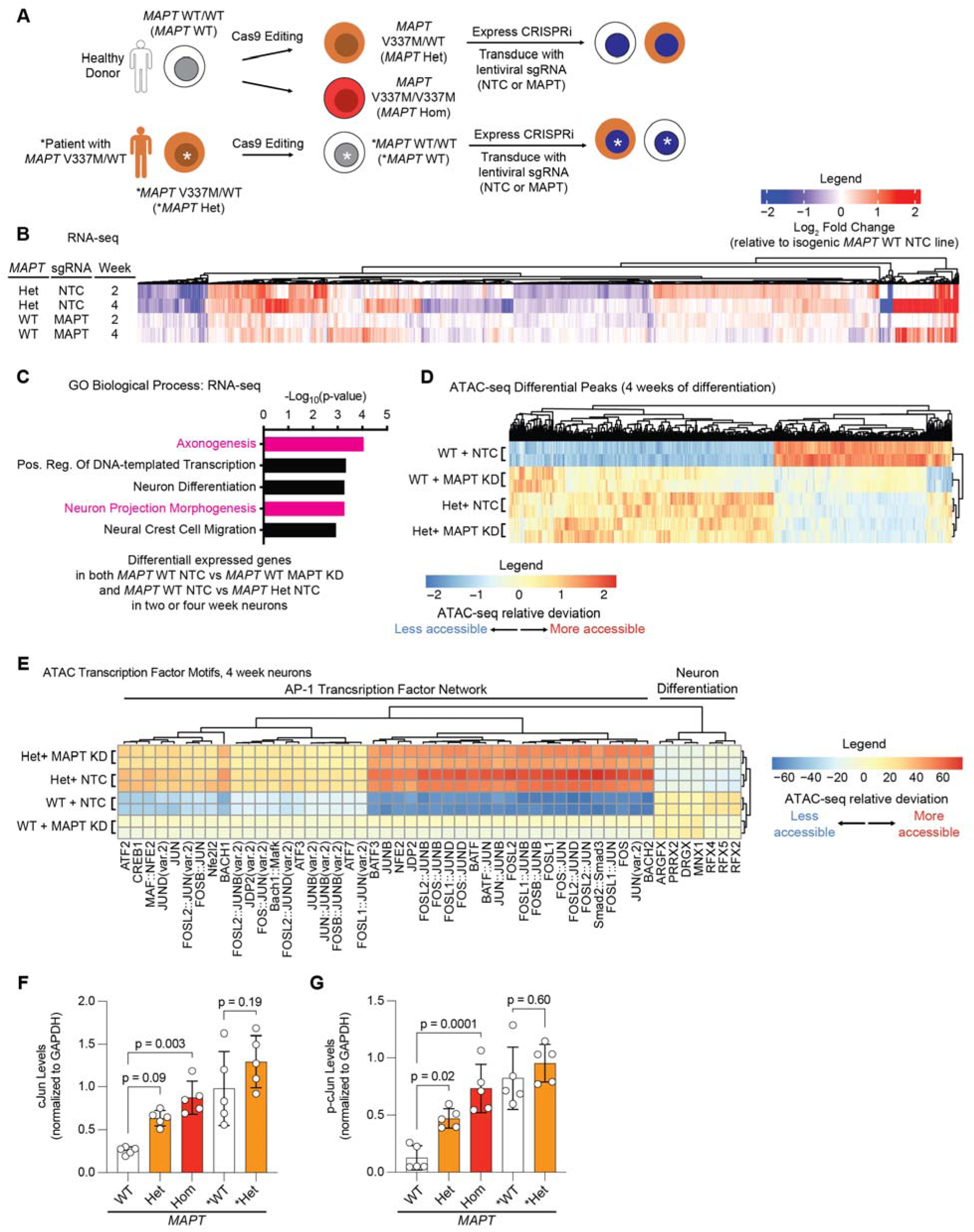
RNA-seq and ATAC-seq in neurons reveal conserved effects of *MAPT* V337M and *MAPT* knockdown on axonogenesis pathways. **(A)** iPSCs from a healthy donor (WTC11, here called *MAPT* WT) or a patient with the heterozygous *MAPT* V337M mutation (GIH6C1, here called \**MAPT* Het) were edited with Cas9 previously to generate a heterozygous *MAPT* V337M clone (*MAPT* Het), a homozygous *MAPT* V337M clone (*MAPT* Hom) and a healthy isogenic control (GIH6C1Δ1E11, here called \**MAPT* WT). These cells were engineered to express dox-inducible mNGN2 in AAVS1 and CRISPRi machinery in CLYBL. We transduce the iPSCs with lentivirus for sgRNA/BFP expression. **(B)** Heatmap comparing changes in gene expression based on RNA-seq in *MAPT* Het NTC and *MAPT* WT MAPT KD vs. *MAPT* WT NTC at 2 and 4 weeks post differentiation. Three independent wells of neurons for each genotype/sgRNA combination were harvested at each timepoint. **(C)** Gene Ontology (GO) term enrichment analysis of the RNA-seq experiment in (B). Genes that are differentially expressed in both *MAPT* Het and *MAPT* WT *MAPT* KD vs. *MAPT* WT NTC were analyzed with Enrichr, and top terms were plotted. Pathways related to axonogenesis and neuron morphology are colored magenta. **(D)** Heatmap summarizing ATAC-seq differential peaks at 4 weeks of differentiation. Two independent wells of neurons for each genotype/sgRNA combination were harvested. **(E)** Heatmap summarizing ATAC-seq transcription factor motif analysis at 4 weeks of differentiation from the same experiment in (D). Clusters were analyzed for pathway enrichment using Enrichr, and major pathways are annotated (“AP-1 Transcription Factor Network” and “Neuron Differentiation”). **(F-G)** Quantification of cJun (F) and p-cJun (G) from the western blots in (S2D). Significance was calculated using one-way ANOVA with Šidák’s multiple comparison test, and comparisons were restricted within the donor background.

RNA-seq of neurons harvested at 2 and 4 weeks of differentiation revealed overlap between effects in *MAPT* Het neurons and *MAPT* WT tau knockdown neurons (Figure 1B). Genes that were differentially expressed in *MAPT* Het neurons and *MAPT* WT tau knockdown neurons were significantly enriched for regulators of axonogenesis (Figure 1C). Knocking down tau in *MAPT* Het neurons resulted in only five differentially expressed genes (Figure S1D).

Differentially expressed genes in *MAPT* Hom and \**MAPT* Het neurons compared to isogenic controls were also significantly enriched for regulators of axonogenesis, even at one week of differentiation (Figure S1E-G). While many of the same transcripts relevant to axonogenesis are perturbed in \**MAPT* Het and *MAPT* Het, we did not see a high level of concordance in direction of change (Figure S1E). On the other hand, the changes in *MAPT* Het and *MAPT* Hom are extremely similar, suggesting high concordance between distinct clones in the same genetic background.

ATAC-seq at 2 and 4 weeks of differentiation showed similar patterns as the RNA-seq (Figure 1D, Figure S2A), and genes with differentially accessible peaks proximal to their transcription start site (TSS) were enriched for axon-related genes (Figure S2B). Transcription factor motif analysis showed that motifs for the AP-1 Transcription factor network, which includes the cJun family of transcription factors, were consistently more accessible in *MAPT* Het and *MAPT* WT tau knockdown neurons compared to controls (Figure 1E, Figure S2C).

Supporting the validity of the ATAC-seq results, we found that both p-cJun and cJun are increased in *MAPT* Het, *MAPT* Hom and *\*MAPT* Het neurons vs. isogenic controls (Figure 1F-G, Figure S2D). The increase in *\*MAPT* Het vs. *MAPT* WT neurons was not statistically significant, likely due to basal cJun activation in control neurons. *MAPT* V337M and tau knockdown induce overlapping changes in chromatin accessibility and transcription of axonogenesis-related genes, suggesting that some early phenotypes in *MAPT* V337M neurons are relevant to normal tau function.

### MAPT V337M and tau knockdown perturb phosphorylation of axonogenesis-related proteins

We hypothesized that changes in cJun and p-Jun may reflect broad changes in intracellular signaling caused by V337M tau. To identify shifts in signaling occurring at early stages of axonogenesis, we determined the total proteome and phosphoproteome of 1-week neurons with *MAPT* V337M and/or tau knockdown by mass spectrometry. Phosphoproteomic analysis of *MAPT* V337M neurons confirmed elevated p-cJun levels while also uncovering differential phosphorylation of proteins regulating neuron projection development and splicing (Figure 2A,C). There was significant overlap in the proteins with differential phosphorylation between *MAPT* Hom, *MAPT* Het and *\*MAPT* Het neurons vs. isogenic controls (Figure 2B), though the identities of the differential phosphosites varied between conditions (Figure S3A). Gene set enrichment analysis for the 56 conserved proteins with changes in phosphorylation in *MAPT* V337M neurons showed that the top enriched terms were related to neuron projection development (Figure 2C). The total protein levels for many of these factors were not significantly changed, suggesting that these changes are due to specific signaling events altering phosphorylation patterns, rather than just changes in protein levels (Figure S3B-D, Figure S9 A-C).

**Figure 2:**
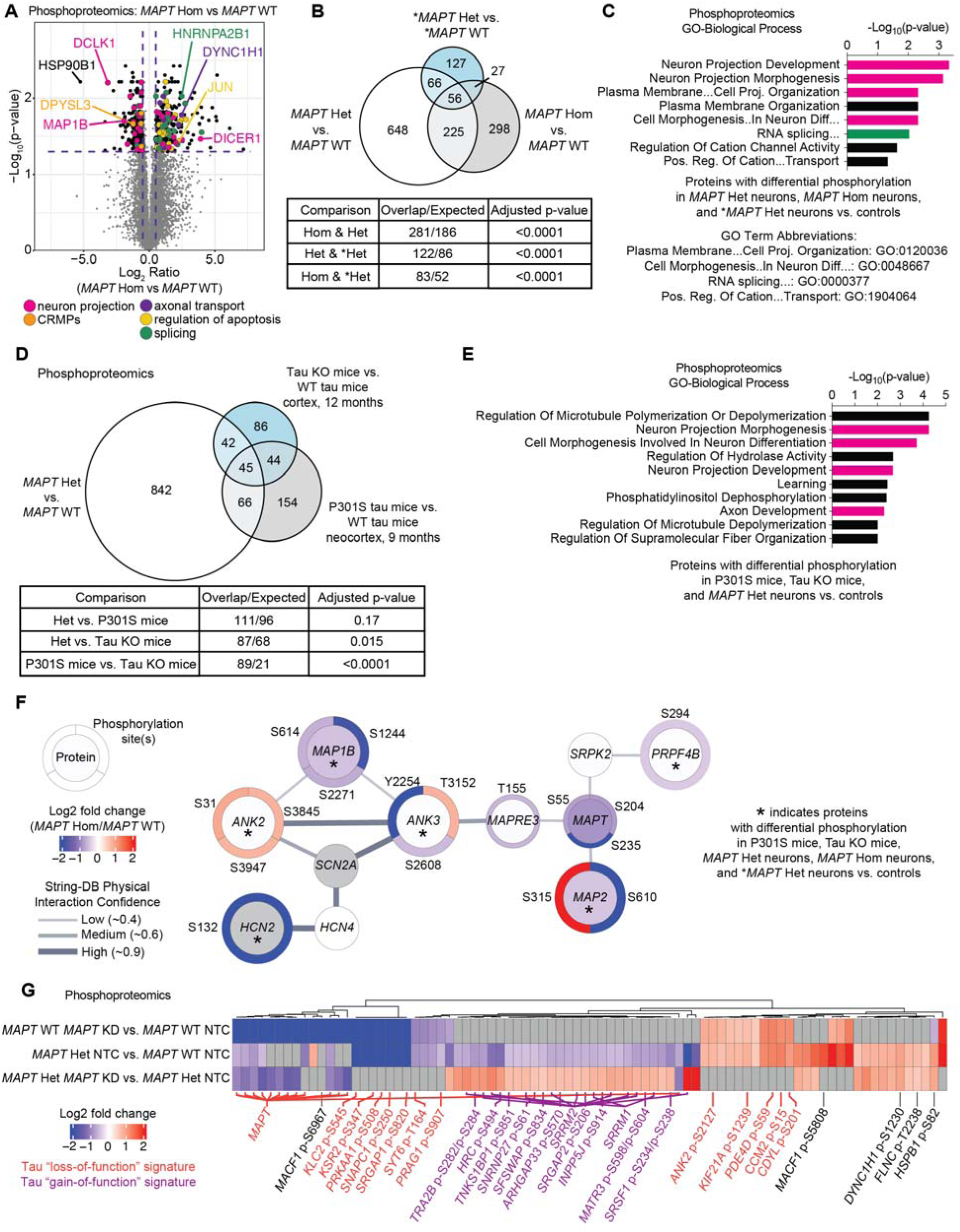
Proteomics uncovers altered phosphorylation of axonogenesis-related proteins in neurons with the *MAPT* V337M mutation. **(A)** Volcano plot showing changes in protein phosphorylation in *MAPT* Hom vs. *MAPT* WT neurons using mass spectrometry. Four independent 150mm dishes of neurons for each condition were harvested after one week of differentiation. Dots represent individual phosphorylation sites. **(B)** Proteins with differential phosphorylation between *MAPT* Het, *MAPT* Hom neurons, and \**MAPT* Het vs. controls. Significance was calculated using multiple t-tests adjusted with Šidák single-step correction. Proteins with differential phosphorylation in all three datasets were filtered to identify 56 conserved proteins. **(C)** GO term enrichment of the 56 proteins from (B). Neuron morphology term bars are magenta, and splicing term bars are green. Top non-overlapping significant terms are shown. **(D)** Proteins with differential phosphorylation between *MAPT* Het vs. *MAPT* WT and two published mouse phosphoproteomics datasets, including tau KO mice and P301S mice vs. WT mice. Significance was calculated using multiple t-tests adjusted with Šidák single-step correction. Proteins with differential phosphorylation in all three datasets were filtered to identify 45 conserved proteins. **(E)** GO term enrichment of the 45 proteins from (D). Annotations are consistent with (C). **(F)** String-DB protein-protein interaction network of proteins with differential phosphorylation in five datasets: *MAPT* Het, *MAPT* Hom, \**MAPT* Het vs isogenic controls, and tau KO mice and P301S mice vs. controls. The inner circle is colored based on the protein log_2_ fold change, and the outer circles are colored based on the log_2_ fold change for the indicated phosphorylation site. **(G)** Heatmap of phosphoproteomics data comparing *MAPT* KD vs. isogenic controls. Phosphosites that are decreased in *MAPT* Het vs. *MAPT* WT but that are rescued by tau knockdown in *MAPT* Het neurons are labeled in purple as a tau “gain-of-function” signature. Phosphosites that are changed in the same direction in *MAPT* WT *MAPT* KD and *MAPT* Het vs. WT are labeled in orange as a tau “loss-of-function” signature.

We next compared our phosphoproteomic datasets to recently published mouse phosphoproteomic datasets using tau knockout mice [24] or P301S tau mice [25]. We found statistically significant overlap for proteins with differential phosphorylation in our data and the tau knockout mice (p = 0.015) but not with the P301S mice (p=0.17) (Figure 2D). However, we noted that there was significant overlap between the tau knockout mice and the P301S mice (p<0.0001). Gene set enrichment analysis identified substantial enrichment of axonogenesis-related protein phosphorylation changes in the 45 conserved proteins with differential phosphorylation in *MAPT* Hom neurons, tau knockout mice, and P301S tau mice (Figure 2E). When we determined an even more focused set of proteins that also have differential phosphorylation in *MAPT* Hom and \**MAPT* Het neurons from patient iPSCs, we found a core network of proteins in highly related pathways regulating neuron morphogenesis and polarity (Figure 2F), including *ANK3* and *MAPRE3*. *ANK3* and *MAPRE3* were recently identified to be important for V337M tau-induced defects in axon initial segment plasticity [17].

We observed two patterns of protein phosphorylation changes due to tau knockdown (Figure 2G). We named phosphorylation changes in *MAPT* Het neurons that phenocopy *MAPT* WT *MAPT* KD the “Tau loss of function” signature. The “Tau gain of function” signature is defined as phosphorylation changes in *MAPT* Het neurons that are rescued by tau knockdown in *MAPT* Het neurons. Many phosphorylation changes were specific to either the *MAPT* Het neurons or tau knockdown in the *MAPT* WT neurons. When we performed gene set enrichment analysis on proteins with differential phosphorylation in *MAPT* WT tau knockdown neurons, the only significantly enriched term was “Regulation of microtubule-based process,” with many of these proteins being involved in axonogenesis (Figure S3E). Gene set enrichment analysis of proteins with differential phosphorylation in *MAPT* Het tau knockdown neurons compared to *MAPT* Het showed that splicing factors were predominantly affected, whereas cytoskeletal and axonogenesis proteins were not perturbed (Figure S3F).

### MAPT V337M and tau knockdown disrupt axonogenesis

To directly test if the *MAPT* V337M mutation causes tau loss of function in axonogenesis, we longitudinally imaged neurons expressing a membrane targeted Lck-mNeonGreen and a nuclear localized NLS-mTagBFP to directly measure axonogenesis (Figure 3A). Intriguingly, *MAPT* KD and the *MAPT* V337M mutation both caused decreased main axon and total neurite length (Figure 3B-D) without significantly perturbing axon branch or secondary neurite length (Figure 3E,F). We validated these results in the patient-derived neurons and found a consistent axonogenesis defect (Figure S4A-C).

**Figure 3:**
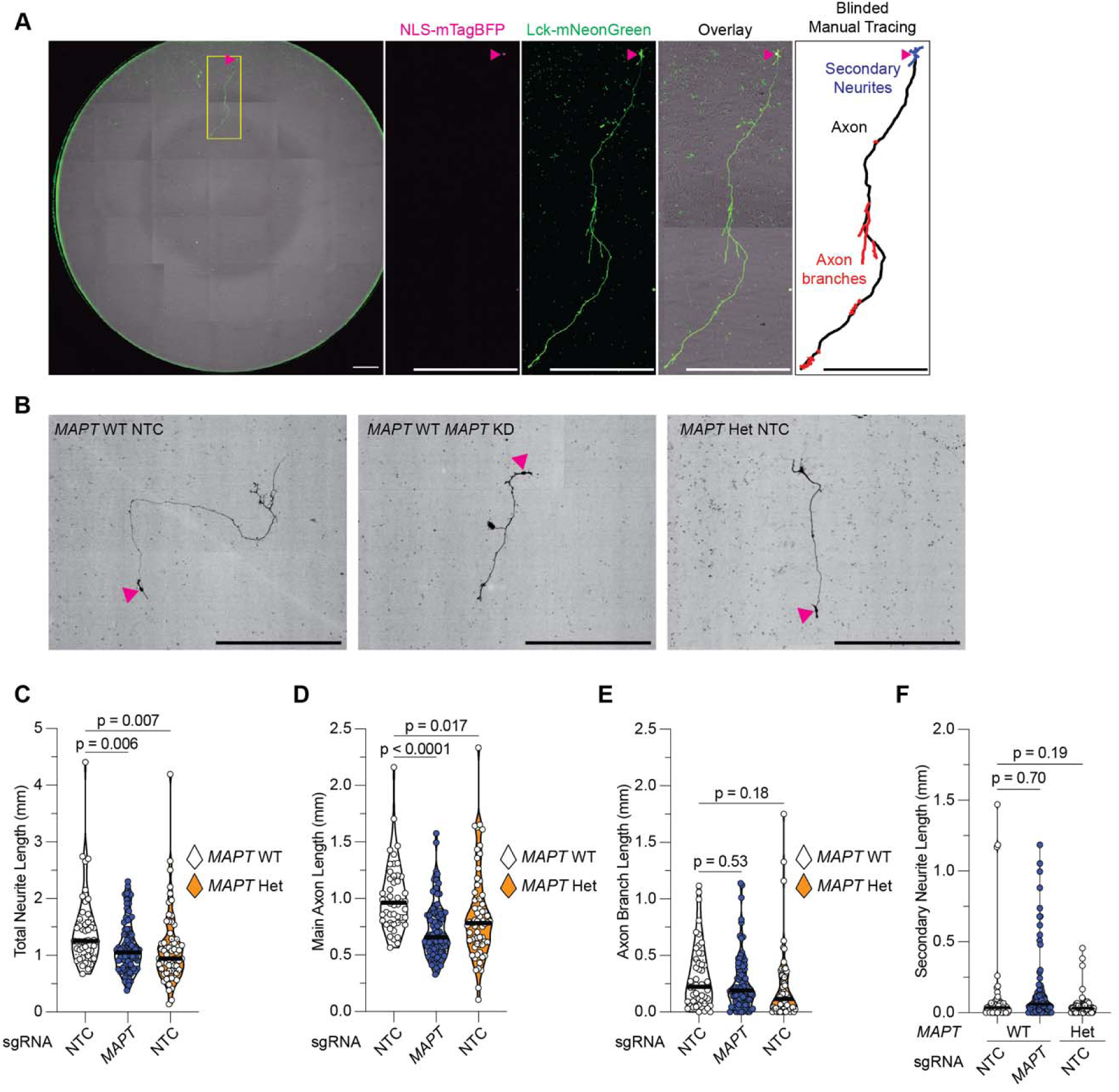
*MAPT* V337M and *MAPT* knockdown disrupt axonogenesis. **(A)** *(From left)* Entire well of a 96 well plate stitched together and overlayed. The nucleus is marked with a magenta arrowhead, and the inset region of interest (ROI) is marked with a yellow box. The scale bars are 500µm. NLS-mTagBFP (magenta) marks the nucleus (magenta arrowhead). Lck-mNeonGreen (green) labels the soma, axon and secondary neurites. Overlay showing the sparse labeled neuron with many unlabeled neurons close by. Example inset of a manually traced neuron showing the features measured. **(B)** Representative images of neurons expressing Lck-mNeonGreen (black). Nuclei are marked with magenta arrowheads, and the scale bars are 200µm. **(C-F)** Total neurite length (C), main axon length (D), axon branch length (E), and secondary neurite length (F) were quantified in day 5 neurons. Significance was calculated using one-way ANOVA with Šidák’s multiple comparison test.

### V337M tau is hypophosphorylated during early differentiation in neurons

We observed that *MAPT* V337M neurons had lower tau phosphorylation compared to WT across all domains of the protein at many sites (Figure 4A and 4B) and validated these changes by western blot (Figure 4C, Figure S5D-E). Many of the differential phosphorylation sites are known to be hyperphosphorylated in Alzheimer’s disease and other tauopathies [27–29] (Figure S5A). Intriguingly, V337M tau hypophosphorylation is transient, approaching WT levels after two to four weeks of differentiation (Figure 4D, Figure S5F-I).

**Figure 4:**
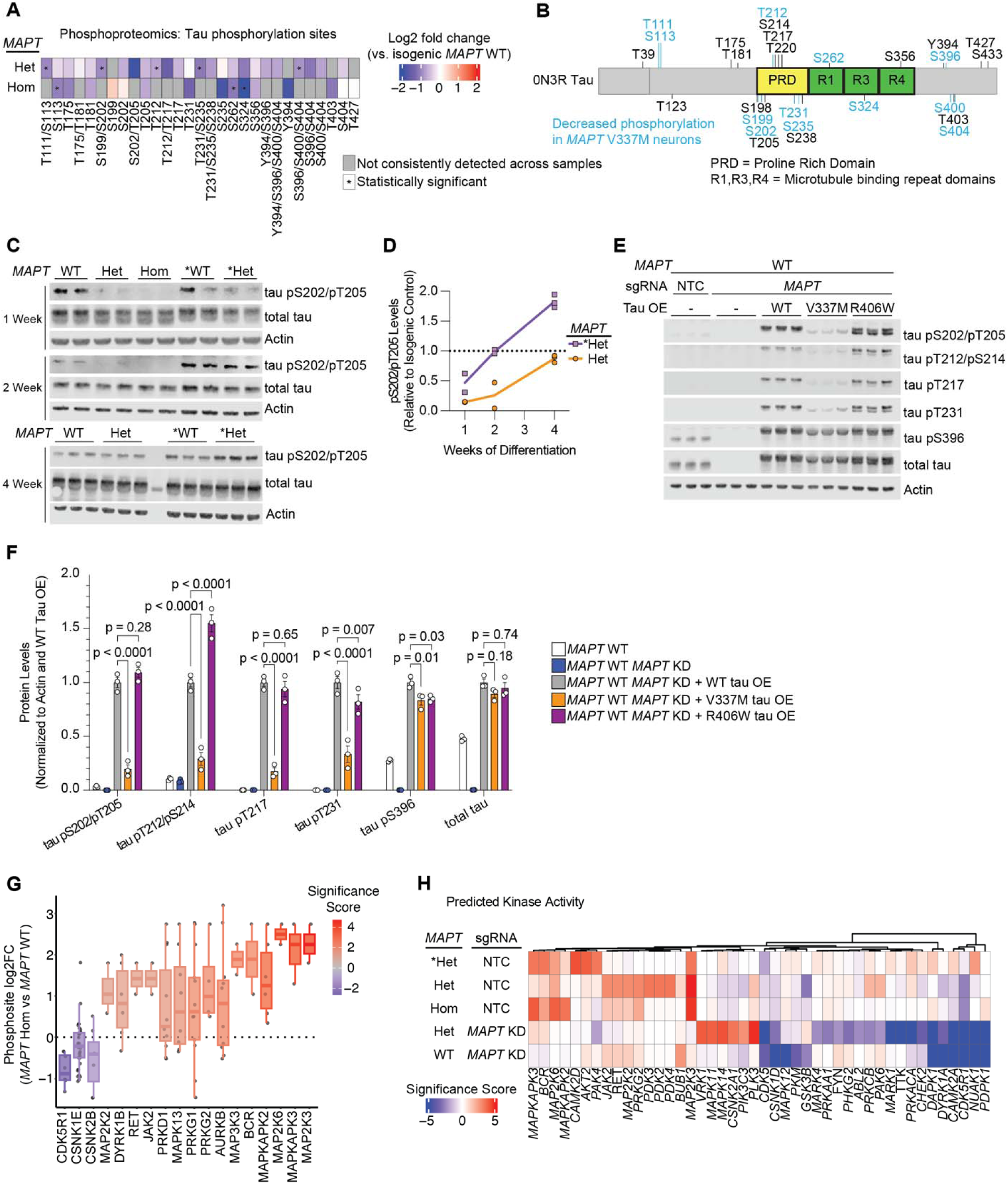
Tau phosphorylation is reduced in neurons with the *MAPT* V337M mutation. **(A)** Heatmap of tau phosphorylation from *MAPT* Hom or *MAPT* Het vs. *MAPT* WT. Phosphosites that were not detected in more than half of the replicates in both samples are marked in grey, and statistically significant phosphorylation changes are marked with an asterisk. **(B)** Protein domain map of 0N3R tau with detected phosphorylation sites labeled. Decreased phosphorylations detected in either *MAPT* Hom or *MAPT* Het neurons are labeled in blue. When phosphoproteomics could not distinguish between multiple potential phosphosites, all are included. **(C)** Western blot validating decreased tau phosphorylation in neurons with V337M tau after one week of differentiation. Two or three independent wells of neurons were harvested after one, two or three weeks of differentiation. AT8 was used to label tau pS202/pT205, and Tau13 was used to label total tau. **(D)** Quantification pS202/pT205 levels normalized to isogenic controls for *MAPT* Het and \**MAPT* Het from the western blots in (C). **(E)** WT, V337M and R406W tau were overexpressed via lentivirus in *MAPT* WT *MAPT* KD iPSCs. Three independent wells of neurons were harvested after one week of differentiation. pTau and total tau levels were analyzed by Western blot. **(F)** Quantification of the western blot in (E). Band intensities were normalized to actin and to the WT tau overexpression line. Significance was calculated using two-way ANOVA with Dunnet’s multiple comparisons test. **(G)** Kinase activity analysis from phosphorylation changes in neurons after one week of differentiation with the homozygous *MAPT* V337M mutation (*MAPT* Hom) vs. isogenic controls (*MAPT* WT). The log_2_ fold change of phosphopeptide abundance for annotated kinase substrates is plotted. The range is represented by the thin lines, the box represents the IQR, and the median is represented by a thick line. **(H)** Heatmap for kinase activity scores from all five phosphoproteomic datasets vs. isogenic controls.

To further explore how V337M tau may have decreased phosphorylation in neurons, we overexpressed WT tau, V337M tau or R406W tau in *MAPT* WT neurons with endogenous tau knocked down. Consistent with our phosphoproteomics results, V337M tau had decreased phosphorylation at numerous sites despite having similar tau levels to WT tau and R406W tau (Figure 4E-F). Intriguingly, R406W tau only had decreased phosphorylation at some of these sites. V337M tau consistently had lower pS202/T205 tau levels when compared to WT or R406W tau at two and four weeks of differentiation when overexpressed (Figure S6A-C). To test if the V337M mutation directly impaired susceptibility to phosphorylation, we purified WT and V337M tau and tested phosphorylation by GSK3B and PKA *in vitro* (Figure S6D-G). The V337M mutation had no significant effect on phosphorylation by GSK3B or PKA for the sites tested. These data suggest that tau variants affect tau phosphorylation in neurons via indirect mechanisms in the cellular context.

Extensive work has been done to characterize tau phosphorylation sites and map them to their kinases [30–35]. Proline-directed phosphorylation sites were decreased in *MAPT* V337M neurons, many of which serve as priming sites for additional sites of decreased tau phosphorylation (Figure S5B,C). Leveraging our global view of phosphorylation changes in *MAPT* V337M neurons, we predicted which kinases may have changes in activity based on known kinase-substrate relationships (Figure 4G,H). Kinases in the p38 MAPK pathway such as *MAP2K3* and *MAP2K6* were predicted to have increased activity in *MAPT* V337M neurons (Figure 4H). *MAPK11* and *MAPK14* targets had increased phosphorylation specifically in *MAPT* V337M neurons with tau knockdown, whereas *MAPK12* substrates had decreased phosphorylation specifically in *MAPT* WT neurons with tau knockdown. Known tau kinases with well-documented roles in tauopathy were also predicted to have differential activity, including *GSK3B*, *CDK5,* and *CDK5R1*. CDK5 and p38 MAPKs are both proline-directed kinases that are known to phosphorylate tau at several sites that had decreased phosphorylation in *MAPT* V337M neurons.

### CRISPR screens uncover regulators of tau phosphorylation in neurons

To directly test which kinases perturb tau phosphorylation in *MAPT* WT and *MAPT* V337M neurons, we employed CRISPRi and CRISPRa screens to test the effects of gene knockdown or overexpression on tau phosphorylation using the AT8 antibody, which detects the tau pS202/pT205 phosphoepitope (Figure 5A, Figure S7A-D). We transduced iPSCs with a lentiviral sgRNA library targeting 2,325 genes encoding kinases, phosphatases and other proteins in the “druggable genome”[36]. Two weeks after differentiation, neurons were fixed and stained with AT8 and sorted based on AT8 signal. Next generation sequencing identified genes that causally regulate AT8 levels. We filtered out hits for enrichment analysis that also modified T22 levels in previously published work (Figure S7E)[37]. Cytoskeleton genes and genes involved in neuron projection development modified tau phosphorylation in both *MAPT* WT and *MAPT* V337M neurons (Figure 4A,B) without altering T22 levels (Figure S7E). Intriguingly, several kinases in the p38 MAPK pathway altered tau phosphorylation specifically in *MAPT* V337M neurons (Figure S8A). Other kinases predicted to have differential activity that may have regulated tau phosphorylation in *MAPT* V337M neurons did not affect tau pS202/pT205 levels, including *CDK5*, *CDK5R1*, and *GSK3B* (Figure S7F). We mapped the detected tau phosphorylation sites in our neurons to their known kinases based on the literature, overlaying phosphorylation sites that were differential in *MAPT* V337M (blue) with kinases whose knockdown or overexpression modified tau phosphorylation at S202/T205 (red) (Figure 5D).

**Figure 5:**
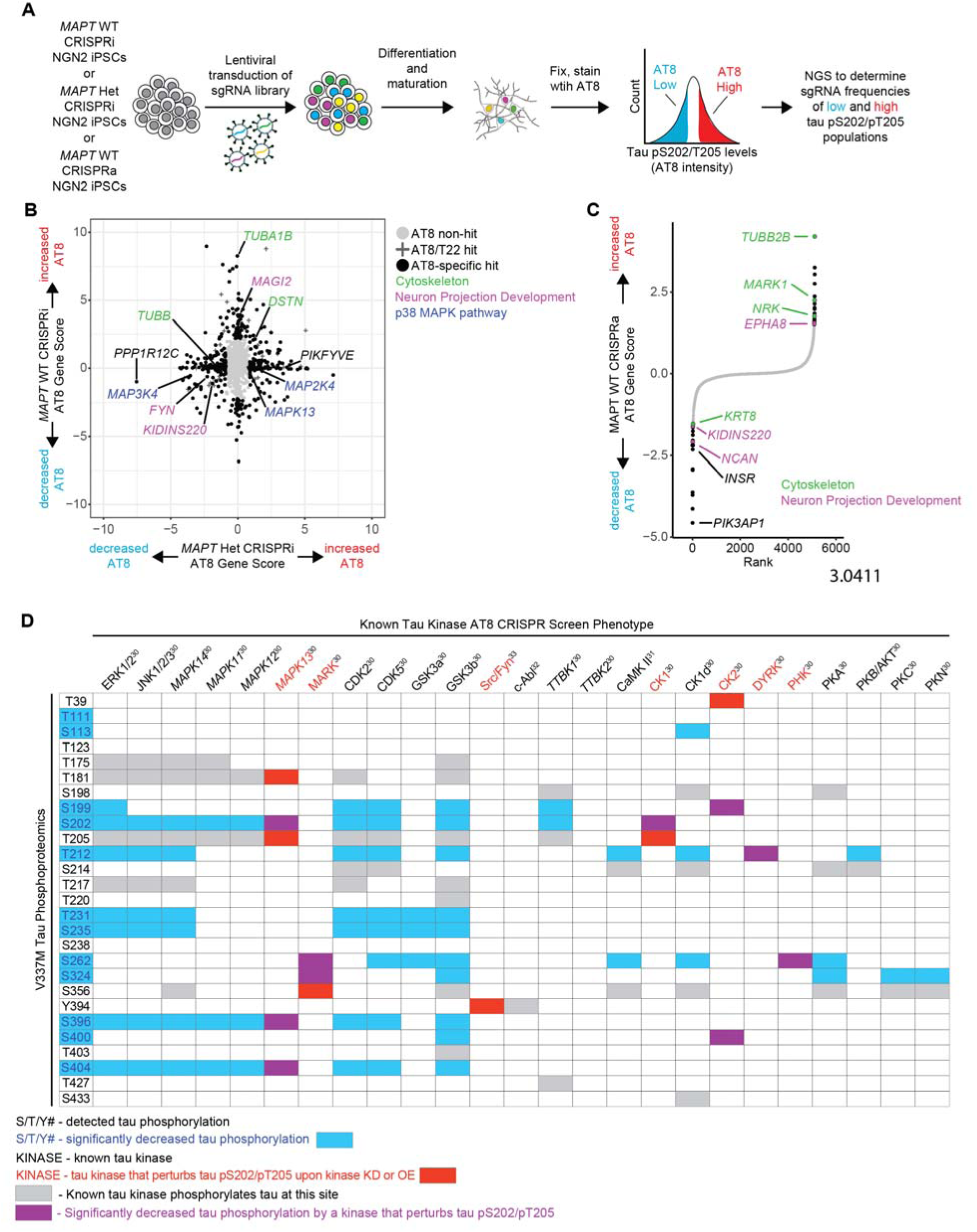
CRISPR screens elucidate regulators of tau phosphorylation in neurons. (A) Pooled genetic screening workflow for tau pS202/pT205 levels in neurons. *MAPT* WT CRISPRi NGN2 iPSCs, *MAPT* Het CRISPRi NGN2 iPSCs or *MAPT* WT CRISPRa NGN2 iPSCs were transduced by lentivirus with a pooled “druggable genome” sgRNA library targeting 2,318 genes enriched for kinases and phosphatases. iPSCs were differentiated into neurons. After two weeks, neurons were fixed, stained with AT8 and sorted for high or low AT8 staining. The high AT8 and low AT8 samples were sequenced to determine which sgRNAs were enriched in either fraction. **(B)** Scatter plot comparing CRISPRi screens in *MAPT* Het neurons vs. *MAPT* WT neurons. AT8 non-hits are labeled with grey circles, AT8 hits that also modify tau levels (using the T22 antibody as a surrogate for total tau levels) are labeled with “+”, and AT8-specific hits are labeled with black circles. Top genotype-specific hits are labeled in black, and key pathways are labeled in green (cytoskeleton), purple (Neuron projection development) and blue (p38 MAPK pathway). **(C)** Rank plot showing the results of the CRISPRa screen in *MAPT* WT neurons. AT8 non-hits are labeled in grey and AT8 hits are labeled in black. Hits that are cytoskeleton-related genes are labeled in green and hits that are related to neuron projection development are labeled in purple. **(D)** Detected tau phosphosites are mapped to the phenotype of known tau kinases from the CRISPRi/a screens. Tau phosphosites that were significantly different in either *MAPT* Hom or *MAPT* Het neurons vs. *MAPT* WT are indicated with blue text/boxes. Kinases whose knockdown or overexpression perturb tau phosphorylation at S202/T205 are indicated by red text/boxes. Overlap between significant kinases and differential phosphorylations are indicated by purple boxes. Grey boxes indicate phosphorylations by known tau kinases that are not significantly differential or involved in tau pS202/T205. References for known tau kinase activity are indicated by the kinase name.

The overlap between differential tau phosphorylation and kinases that regulate pS202/pT205 in neurons (purple) narrows the list down to a few candidate kinases. Tau phosphorylation at different sites and by different kinases has been shown to be highly interdependent, and p38 MAPK had the highest interdependence of the kinases tested [38]. Another fascinating signature emerged when we looked at proteins involved in arginine/citrulline/nitric oxide metabolism that were very consistently differentially expressed (Figure S8A). Nitric oxide activates guanylate cyclases, which in turn activate cGMP-dependent kinases [39]. *PRKG1* knockdown increased AT8 levels specifically in *MAPT* Het neurons (Figure S8A). Overexpression of *MARK1*, a kinase that phosphorylates tau in the microtubule binding domain and regulates tau’s interaction with microtubules, caused increased tau phosphorylation in *MAPT* WT neurons (Figure 5C). This is consistent with previous work showing that phosphorylation at S262, S324 and S356 affects phosphorylation sites distal from the microtubule binding domain [40].

## Discussion

We have discovered that an FTD-causing variant of tau leads to early tau hypophosphorylation and perturbs axonogenesis pathways in differentiating neurons, overlapping at least in part with effects seen in tau knockdown. These findings are surprising because disease-associated tau is typically associated with increased tau phosphorylation and would not be expected to have shared phenotypic overlap with tau loss. Other groups have shown in mice or in primary neurons that reducing tau can have varying effects on axonogenesis. Acute tau ablation in mouse neurons *in vitro* prevents axonogenesis by inhibiting polarization [41, 42] and tau knockout in primary hippocampal neurons and human iPSC-derived neurons reduces neurite outgrowth [43–45]. hTau overexpression rescued neurite outgrowth defects in tau knockout neurons, and phosphorylation at tau T205 was necessary to rescue this defect. [43] Tau phosphorylation was shown to temporarily increase during development in rats during the period of active neurite outgrowth [46].

We found that there were changes in axon-related genes at the RNA, chromatin accessibility, and phosphoproteomic level without obvious changes at the protein level. There are two possible explanations for this discrepancy. RNA-seq and ATAC-seq are much more sensitive than proteomics, so many of the differentially expressed genes are not detected in the proteomics datasets. At the same time, the axon-related proteins that are differentially phosphorylated do not have strong RNA-seq or total proteomics changes, suggesting that these are truly changes at the phosphoprotein level specifically. One could image a mechanism where tau perturbs axonogenesis, leading to rapid phosphorylation signaling changes and later transcriptomic changes to compensate for the deficit.

We acknowledge that there are limitations to our study. Our neurons under the conditions we used only express a single isoform of tau, the fetal isoform 0N3R. Understanding how different tau isoforms are regulated and how they contribute tau function in health and disease is an open question. Additionally, it will be intriguing to understand how different disease variants of tau perturb neurons. Our data showing phosphorylation differences between WT tau, V337M tau and R406W tau joins a growing body of literature showing that different mutations have different effects on tau properties, including microtubule binding, microtubule polymerization, and fibril formation [47–53]. Our data highlights a link between the V337M mutation and tau loss of function at an early timepoint, but we do not have mechanistic insight into how the mutation causes tau loss of function. A major open question from this work is how the V337M mutation leads to lower tau phosphorylation and changes in phosphorylation of other axonogenesis proteins. Understanding how tau perturbations interact with other key regulators of microtubule dynamics like *MAP1B* and *MAP2* will give new insights on the physiological regulation of these proteins and the cellular responses to tau loss or mutation. Our results from our phospho-tau CRISPR screens overlaid with the V337M tau phosphoproteomics data give interesting clues into which tau regulators might be responsible for V337M tau hypophosphorylation. The p38 MAPK pathway, MARK, CK1, CK2, PKG, DYRK and PHK are all candidates that could be involved tau phosphorylation during development or the cellular response to mutant tau during axonogenesis. Tau phosphorylation by p38 MAPK was shown to be highly interdependent, and p38 MAPK was necessary for neurite outgrowth in PC12 cells [38, 54]. Taken together, our data strongly supports the role of p38 MAPKs in regulating axonogenesis and tau phosphorylation. Additional mechanistic work and characterization in other model systems will help the field better understand the physiological regulation of tau in development and how disease variants of tau perturb development.

The question remains whether a loss of tau function would have adverse effects to a disease variant carrier throughout life. The precise physiological roles of tau are unclear and have been debated for many years [4, 22]. This is in large part due to the many conflicting studies, both in physiological and pathogenic contexts. Given the earlier results in showing the importance of tau for axonogenesis, it was expected that knocking out tau in mice would be lethal and that tau would be essential for neurodevelopment. Early mouse studies showed that tau knockout was surprisingly well tolerated [55]. There were no obvious defects in polarization or gross morphology, but microtubules in small caliber axons were destabilized. *Map1a* was upregulated in tau knockout mice, suggesting that the mice were compensating for tau loss. This could explain the difference in phenotypes as compared to the acute depletion of tau with ASOs. Knocking out tau and *Map1b,* another microtubule-associated protein, leads to much more severe phenotypes than either knockout individually [56]. Dawson et al. disputed the findings of Harada et al. due to poor WT data [57]. In their work, they found that indeed tau knockout did cause a delay in neurite outgrowth and axonogenesis.

Biswas and Kalil showed that tau knockout neurons had altered microtubule dynamics in growth cones, resulting in a change in overall growth cone morphology [58]. Microtubules were less bundled, and microtubule polymerization directionality as measured by EB3 was more dispersed in tau knockout neurons. There also was a reduction in tyrosinated tubulin projecting into the filopodia of the peripheral domain. Another paper showed that tau knockout increased Fyn mobility in dendrites and lowered Fyn localization in dendrites and spines [59]. Intriguingly, expressing P301L tau had the opposite effect and anchored or trapped Fyn in dendritic spines.

Many motor and behavioral phenotypes have been observed in tau knockout mice. Tau knockout mice or mice with acute tau reduction with antisense oligonucleotides have consistently shown resistance to seizures [21, 60–63]. Another consistent theme is that there are often behavioral and learning changes in tau knockout mice, including hyperactivity, fear conditioning, and memory [64–69]. There is more controversy over the effect of tau knockout on motor function. Some groups report motor deficits in tau knockout mice [64, 70, 71], while others claim there are no significant changes in tau knockout mice to motor function [60, 61, 69]. One group showed that tau is essential for long term depression in the hippocampus [72], while another showed that tau knockout only perturbs long term potentiation [69]. Tau phosphorylation has also been shown to be required for long term depression [73].

Considering our data in the context of these other findings, we expect that loss-of-function phenotypes would coincide with the onset of tau expression and axon extension. Tau loss of function may precede human disease onset by many decades, occurring during development and continuing through adulthood via perturbed synaptic plasticity. A study showed that mice with the *MAPT* P301L mutation show early cognitive changes before tau pathology is detectable [74]. A Parkinson’s disease GWAS study found that *MAPT* was a significant risk locus for Parkinson’s disease that is uncoupled from the age of onset [75]. Ye and colleagues proposed that tau may drive changes during development or early in life that then increase risk for Parkinson’s disease decades later [76]. Two studies have also identified cognitive differences between *MAPT* carriers and non-carrier siblings decades before expected disease onset [77, 78].

Our work also emphasized the importance of having iPSCs from multiple individuals and multiple clones paired with appropriate controls, such as tau knockdown or knockout. Furthermore, comparisons to other published data sets revealed previously underappreciated relationships, including overlapping molecular phenotypes between *MAPT* knockout and *MAPT* P301S mice. It will be fascinating to uncover the mechanisms of these shared signaling pathway changes and to determine if they are due a shared stress response, or if downstream phenotypes converge despite unique upstream perturbations. Previous work using different differentiation protocols and much longer time scales showed that FTD-causing tau variants cause tau hyperphosphorylation in more mature neurons, suggesting that there is a complex, time-dependent effect of *MAPT* mutations on developmental tau phosphorylation patterns. There was substantial overlap between our RNA-seq findings and a recent paper using *MAPT* V337M neurons in an organoid model, which is interesting because of the observed tau hyperphosphorylation at the later timepoints in their model [9]. Intriguingly, one other group reported decreased tau phosphorylation in organoid-derived iPSC neurons with *MAPT* R406W. [79] Earlier work showed that fetal tau was highly phosphorylated during development, but the precise mechanisms and functions of this process are still unknown.[32, 80–82] Our findings suggest that at earlier timepoints different tau mutations may behave in unexpected ways and have complex effects on cellular pathways. [5, 9, 83]

Beyond the findings presented here, we expect that the data sets we have generated will continue to be useful to the field as we resolve the plethora of molecular and cellular phenotypes driven by pathogenic tau in a variety of contexts. Similarly, although much is still to be learned about the consequences of dysregulated tau phosphorylation (both loss and gain), our functional genomic screens could inform the design of tau phosphorylation modulators, perhaps even for therapy.

## Conclusions

Our study aims to characterize WT, V337M tau and tau knockdown neurons to understand how tau loss or mutation perturbs neuron biology. We show that V337M tau and tau knockdown have conserved effects in RNA-seq, ATAC-seq and phosphoproteomics. Surprisingly, we found that V337M tau causes tau hypophosphorylation. We performed functional genomics screens to uncover the regulators of tau phosphorylation in WT and V337M tau neurons. V337M tau perturbs axon morphology pathways similarly to tau loss and causes tau hypophosphorylation, which could contribute to the previously reported cognitive changes in preclinical *MAPT* variant carriers.

## Materials and Methods

### Human iPSC culture and neuronal differentiation

Human iPSC culture and neuronal differentiation were essentially carried out as previously described [84]. Briefly, human iPSCs from the WTC11 background were cultured in StemFlex Medium (GIBCO/Thermo Fisher Scientific; Cat. No. A3349401). Human iPSCs from the GIH6C1 background were cultured in mTeSR Plus medium (StemCell Technologies; Cat. No. 100-0276). iPSCs were grown in plates or dishes coated with Growth Factor Reduced, Phenol Red-Free, LDEV-Free Matrigel Basement Membrane Matrix (Corning; Cat. No. 356231) diluted 1:100 in Knockout DMEM (GIBCO/Thermo Fisher Scientific; Cat. No. 10829-018). StemFlex Medium was replaced daily. When cells reached 80-90% confluency, cells were dissociated with StemPro Accutase Cell Dissociation Reagent (GIBCO/Thermo Fisher Scientific; Cat. No. A11105-01) at 37°C for 5 min, centrifuged at 200 g for 5 min, resuspended in StemFlex Medium supplemented with 10 nM Y-27632 dihydrochloride ROCK inhibitor (Tocris; Cat. No. 125410) and placed onto Matrigel-coated plates or dishes. Studies at UCSF with human iPSCs were approved by the Human Gamete, Embryo, and Stem Cell Research (GESCR) Committee.

For individual gene knockdown in CRISPRi iPSCs, sgRNAs were introduced into iPSCs via lentiviral delivery. Cells were selected by 1 µg/ml puromycin for 2-4 days and recovered for 2-4 days. Phenotypes were evaluated 5-7 days after infection.

The WTC11 CRISPRi iPSC lines were previously engineered to express mNGN2 under a doxycycline-inducible system in the AAVS1 safe harbor locus. The GIH6C1 iPSC lines were engineered in this work to express Ngn2 under a doxycycline-inducible promoter in the AAVS1 safe harbor locus. For their neuronal differentiation, we followed our previously described protocol [85]. Briefly, iPSCs were pre-differentiated in matrigel-coated plates or dishes in N2 Pre-Differentiation Medium containing the following: Knockout DMEM/F12 (GIBCO/Thermo Fisher Scientific; Cat. No. 12660-012) as the base, 1X MEM Non-Essential Amino Avids (GIBCO/Thermo Fisher Scientific; Cat. No. 17502-048), 10 ng/mL NT-3 (PeproTech; Cat. No. 450-03), 10 ng/mL BDNF (PeproTech; Cat. No. 450-02), 1µg/mL Mouse Laminin (Thermo Fisher Scientific; Cat. No. 23017-015), 10 nM ROCK inhibitor and 2µg/mL doxycycline to induce the expression of Ngn2. After three days, or Day 0, pre-differentiated cells were dissociated with accutase and plated into BioCoat Poly-D-Lysine-coated plates or dishes (Corning; assorted Cat. No.) in Classic N2 neuronal medium or BrainPhys Neuronal Medium. Classic N2 neuronal medium contained the following: half DMEM/F12 (GIBCO/Thermo Fisher Scientific; Cat. No. 11320-033) and half Neurobasal-A (GIBCO/Thermo Fisher Scientific; Cat. No. 10888-022) as the base, 1X MEM Non-Essential Amino Acids, 0.5X GlutaMAX Supplement (GIBCO/Thermo Fisher Scientific; Cat. No. 35050-061), 0.5X N2 Supplement, 0.5X B27 Supplement (GIBCO/Thermo Fisher Scientific; Cat. No. 17504-044), 10 ng/mL NT-3, 10 ng/mL BDNF and 1 µg/mL Mouse Laminin. BrainPhys Neuronal Medium was comprised of the following: BrainPhys Neuronal Medium (StemCell Technologies; Cat. No. 05791) as the base, 0.5x N2 Supplement, 0.5X B27 Supplement, 10 ng/mL NT-3, 10ng/mL BDNF, and 1 µg/mL Mouse Laminin. Neuronal medium was fully changed on day 3 post differentiation and then half-replaced on day 7 and weekly thereafter.

### GIHC1 iPSC cell line generation

GIH6C1 and GIH6C1Δ1E11 were obtained from NeuraCell [23]. iPSCs were transfected with pC13N-dCas9-BFP-KRAB and TALENS targeting the human CLYBL intragenic safe harbor locus (between exons 2 and 3) (pZT-C13-R1 and pZT-C13-L1, Addgene #62196, #62197) [86] using DNA In-Stem (VitaScientific). At the same time, the iPSCs were also transfected with pUCM-AAVS1-TO_hNGN2 (Addgene #105840) [87] and TALENS targeting the human AAVS1 intragenic safe harbor locus (pTALdNC AAVS1_T1, pTALdNC AAVS1_T2). [88] After two weeks, BFP-positive iPSCs (CRISPRi+/mNGN2-), mCherry-positive iPSCs (CRISPRi-/mNGN2+) and BFP/mCherry-positive iPSCs (CRISPRi+/mNGN2+) were isolated via FACS sorting. Cells were plated sparsely in a 10 cm dish (5,000-10,000 per dish) and allowed to grow up until they formed large colonies. Homogenous BFP+/mCherry+ colonies were picked with a pipette tip and placed into a 24 well plate for expansion and characterization. Cre mRNA was then transfected into the iPSCs to remove the selection marker and mCherry. Cells were sorted for mCherry negativity, and then mCherry negative colonies were picked and genotyped.

### Western blots

Neurons were washed 3 times with ice-cold PBS. Ice-cold RIPA with protease and phosphatase inhibitors was added to cells. Lysates were incubated on ice for 2 minutes and then scraped down. Lysates were centrifuged at 12500xg for 10 minutes at 4 °C. The supernatants were collected, and the concentrations were measured with the BCA assay (Thermo Fisher Scientific; Cat No. 23225). 10-20 µg protein were loaded onto 4-12% Bis-Tris polyacrylamide gel (Thermo Fisher Scientific; Cat No. NP0336BOX) Nitrocellulose (BioRad, Cat. No. 1620146) or PVDF membranes were used to transfer the protein in a BioRad Transblot for 11 minutes at 25 V, 25 A. Membranes were then blocked for 1 hour with Licor Intercept blocking buffer (Licor, Cat. No. 927-60001) at room temperature. Primary antibody was added in Licor Intercept block overnight at 4 °C. Blots were washed 3 times for 5 minutes with TBST at room temperature. Secondary antibodies were added in Licor Intercept block for 1 hour at room temperature. Blots were washed 3 times for 5 minutes with TBST at room temperature and imaged on a Licor Odyssey. Immunoblots were quantified by intensity using ImageStudio (Licor).

### Bulk RNA sequencing sample preparation

RNA was harvested from day 7, day 14 and day 28 post differentiation neurons using a Zymo microprep kit (Zymo Research, Cat No. R2062). The library was prepared by first depleting ribosomal RNA (New England BioLabs, Cat No. E7405L). cDNA synthesis was then performed on all remaining RNAs (New England BioLabs, Cat. No. E7765S). Paired-end (PE65) sequencing was performed at the Chan Zuckerberg Biohub and the UCSF Center for Advanced Technology.

### ATAC-seq sample preparation

Omni-ATAC-seq was performed as previously described.[89] In short, nuclei from 50,000 neurons were resuspended with Tn5 transposase (to tag and cleave open chromatin with PCR adapters) and incubated at 37 C for 30 minutes on a thermomixer at 1,000 rpm. DNA was then extracted using the Qiagen MinElute Reaction Cleanup Kit (Cat#28204). Tagged sequences were amplified using Illumina/Nextera i5 common adapter and i7 index adapters. DNA libraries were purified using AMPure XP beads (A63880), and paired-end (PE65) sequencing was performed at the Chan Zuckerberg Biohub and the UCSF Center for Advanced Technology.

### Proteomics sample preparation

Briefly, neurons were scraped off 15 cm dishes at day 7 of differentiation and flash frozen in liquid nitrogen. Cell pellet was lysed by adding 1 ml of 6 M GnHCl, 100mM Tris pH 8 and boiling at 95 C for 5 minutes two times with 5 min rest in between. DNA was sheared three times via probe sonication at 20% amplitude for 10 s., followed by 10 s of rest. Following sonication, samples were allowed to solubilize on ice for 20 mins before clearing cell debris by centrifugation at 16,000 x g for 10 mins and determining protein concentration was using Protein Thermo Scientific 660 assay. Enough lysate for 1 mg of protein was aliquoted and Tris 2-carboxyethyl phosphine (TCEP) and chloroacetamide (CAA) were added to each sample to a final concentration of 40 mM and 10 mM respecitively, before incubating for 10 min at 45 C with shaking. Guanidine was then diluted at least 1:5 with 100 mM Tris pH 8. Trypsin and LysC (Promega) were added at a 1:100 (enzyme:protein w:w) ratio (total protease:protein ratio of 1:50) and digested overnight at 37°C with shaking. Following digestion, 10% trifluoroacetic acid (TFA) was added to each sample to a final pH ∼2. Samples were desalted under vacuum using Sep Pak tC18 cartridges (Waters). Each cartridge was activated with 1 mL 80% acetonitrile (ACN)/0.1% TFA, then equilibrated with 3 × 1 mL of 0.1% TFA. Cartridges were then washed with 4 × 1 mL of 0.1% TFA, and samples were eluted with 0.8 mL 50% ACN/0.25% formic acid (FA). 20 μg of each sample was kept for protein abundance measurements, and the remainder was used for phosphopeptide enrichment. Samples were dried by vacuum centrifugation.

### Phosphopeptide enrichment

For phosphopeptide enrichment of samples for phosphoproteomics, IMAC beads (Fe-IMAC from Cube Biotech) were prepared by washing 3x with washing buffer (0.1% TFA, 80% ACN). Dry, digested peptide samples were resuspended in washing buffer and incubated for 15 mins at 37 C with shaking. Peptides were enriched for phosphorylated peptides using a King Fisher Flex (KFF). A more detailed KFF protocol can be provided. Briefly, after resuspension peptides were mixed with beads and bound peptides were washed three times with wash buffer before being eluted from beads using 50% ACN, 2.5 % NH4OH solution. Enriched phosphorylated peptide samples were acidified using 75% ACN, 10% FA (at a ratio of 5:3 elution buffer:acid buffer), and filtered by centrifugation through NEST tips.

### Mass spectrometry data acquisition

Digested samples were analyzed on an Orbitrap Exploris 480 mass spectrometry system (Thermo Fisher Scientific) equipped with either an Easy nLC 1200 or Neo Vanquish ultra-high pressure liquid chromatography system (Thermo Fisher Scientific) interfaced via a Nanospray Flex source. Separation was performed using a 15 cm long PepSep column with a 150 um inner diameter packed with 1.5um Reprosil C18 particles. Mobile phase A consisted of 0.1% FA, and mobile phase B consisted of 0.1% FA/80% ACN. Abundance samples were separated by an organic gradient from 4% to 30% mobile phase B over 62 minutes followed by an increase to 45% B over 10 minutes, then held at 90% B for 8 minutes at a flow rate of 600 nL/minute.

Phosphoproteomics samples were separated by an organic gradient from 2% to 25% mobile phase B over 62 minutes followed by an increase to 40% B over 10 minutes, then held at 95% B for 8 minutes at a flow rate of 600 nL/minute. To expand the spectral library, two samples from each set of replicates was acquired in a data dependent manner. Data dependent analysis (DDA) was performed by acquiring a full scan over a m/z range of 350-1100 in the Orbitrap at 60,000 resolving power (@200 m/z) with a normalized AGC target of 300%, an RF lens setting of 40%, and a maximum ion injection time of “Auto”. Dynamic exclusion was set to 45 seconds, with a 10 ppm exclusion width setting. Peptides with charge states 2-6 were selected for MS/MS interrogation using higher energy collisional dissociation (HCD), with 20 MS/MS scans per cycle. MS/MS scans were analyzed in the Orbitrap using isolation width of 1.6 m/z, normalized HCD collision energy of 30%, normalized AGC of 200% at a resolving power of 15,000 with a 22 ms maximum ion injection time. Similar settings were used for data dependent analysis of phosphopeptide-enriched and abundance samples. Data-independent analysis (DIA) was performed on all samples. An MS scan at 60,000 resolving power over a scan range of 350-1100 m/z, a normalized AGC target of 300%, an RF lens setting of 40%, and the maximum injection time set to “Auto”, followed by DIA scans using 20 m/z isolation windows over 350-1100 m/z with a 2 m/z overlap at a normalized HCD collision energy of 30%.

### Longitudinal Neurite Imaging

20,000 neurons were plated in Black Corning Biocoat clear bottom 96-well plates (Corning #356640). The background cells were unlabeled *MAPT* WT neurons without any markers or sgRNAs. Neurons labeled with LCK-mNeonGreen were plated at a density of 2-10 cells per well and were imaged every other day from day 1 to day 11 of differentiation. Images were collected on an InCell6000 using a 10x objective. Thirty-six fields per well with 15% overlap were used to image the entire well.

### Purification of recombinant tau protein

Purification of WT 0N4R and MAPT V337M 0N4R recombinant tau protein was performed as previously described [37] with some modifications. Briefly, WT 0N4R tau or V337M 0N4R tau DNA sequences were amplified using primers designed for Gibson DNA assembly (Supplementary Table S6) into the His-SUMO pAJS1085 vector cut with BsaI. Cloned constructs were transformed into BL21 Rosetta2 DE3 cells (Novagen; Cat No.70954-3). 100 mL of overnight LB starter culture was used to inoculate 1 L of LB that was grown at 37°C to O.D of 1.0. Cells were induced with 1mM IPTG (Teknova; Cat No. I3430) for 4 hours at 37 °C then spun down at 4,000xg for 15 minutes at 4 °C. Supernatant was discarded and the pellet was resuspended in 50mL of lysis buffer (150 mM NaCl, 25 mM HEPES pH 7.5, 50 mM imidazole pH 7.5, 2 mM TCEP) supplemented with protease inhibitors (Roche; Cat No.11697498001). Cells were lysed by sonication (50% amplitude with intervals of 5s on and 10s off for a total of 7 minutes on) on ice and then spun at 40,000xg for 30 minutes at 4 °C. The supernatant was loaded onto a nickel affinity column (HisTrap FF, Cytiva; Cat No. 17528601) and eluted with a linear gradient of 50 mM to 500 mM imidazole. The eluted fractions were run on a gel, pooled and then dialyzed overnight at 4 °C with 0.25 mg ULP1 protease into low salt buffer (50 mM MES pH 6, 0.5 mM EDTA, 20 mM NaCl, 2 mM TCEP). This sample was loaded onto a cation exchange column (HIPrep SP HP, Cytiva; Cat No. 29018183) and eluted with a linear gradient of 0-100% high salt buffer (50 mM MES pH 6, 0.5 mM EDTA, 1M NaCl, 2 mM TCEP). Fractions containing tau were pooled and concentrated with column concentrators (Vivaspin protein concentrator, 10,000 MWCO; Cat No. 28932360) and then run on a size exclusion column (HiLoad Superdex 200, Cytiva; Cat No. 28989335). Fractions containing purified tau were pooled, concentrated and dialyzed into tau storage buffer (10 mM phosphate buffer pH 7.2, 1 mM TCEP).

### In-vitro phosphorylation assay

WT 0N4R tau or *MAPT* V337M 0N4R tau (50 µM) were incubated in 30 µl of phosphorylation buffer (25 mM HEPES, 50 mM NaCl, 2 mM DTT, 5 mM MgCl2, 2 mM ATP, 1 mM PMSF) with either 0.4 µg GSK3β (Acro Biosystems; Cat No. GSB-H5545) or 2 µg cAMP-dependent PKA (New England Biolabs; Cat No. P6000S) for 24 hours at 30 °C, shaking at 250 rpm on the Eppendorf ThermoMixer® C using a heated lid. 15 µl of samples were used for western blotting with phospho-tau-specific antibodies, as described in *Western blots*.

### Antibodies used in this study

cJun (CST, #9165)

p-cJun (CST, #91952)

Tau13 (Santa Cruz Biotechnology, sc-21796)

AT8 (Invitrogen, MN1020)

AT100 (Invitrogen, MN1060)

AT180 (Invitrogen, MN1040)

Tau pT217 (Invitrogen, 44-744)

Tau pS396 (Invitrogen, 44-752G)

GAPDH (Santa Cruz Biotechnology, sc-47724)

β-Actin (CST, #4967)

### Molecular Cloning

Overexpression constructs were generated using our previously described PSAP expression vector [84] as a backbone. This vector expressed PSAP fused to a c-terminal mScarlett. We cloned emGFP-BRD2 into this vector (deleting PSAP-mScarlett) and then used XhoI and AgeI restriction enzyme sites to clone in 0N3R tau. We then cloned a gene block for ORF-BamHI-(GS)4-exFlag-T2A-mApple into the vector using the AgeI and EcoRI sites [90]. We then mutated WT 0N3R tau to V337M and R406W to generate the final overexpression constructs.

### CRISPR screening

45 million iPSCs were infected with lentivirus encoding for the H1 sublibrary (Horlbeck et al Elife) at an MOI of ∼0.3 and selected with 1ug/mL puromycin until 100% BFP positive.

Lentivirus preparation was performed as described (https://dx.doi.org/10.17504/protocols.io.8dfhs3n, [90], [85]). For CRISPRa screens, TMP was added at a final concentration of 50uM for all cultures after selection. Cells were then differentiated and cultured as previously described (http://10.17504/protocols.io.bcrjiv4n, [85]). Upon differentiation, pre-differentiated cells were plated on three 15cm PDL-coated dishes at a density of 15 million cells per plate. Neurons were then matured for two weeks. At two weeks of age, neurons were lifted with papain and zinc fixed as previously described [90]. On the day of sorting, preparation for FACS was performed as described [90] using the AT8 antibody (Thermo MN1020) at a concentration of 1:200. After sorting, cells were pelleted at 200xg for 20 minutes, the supernatant was removed and the pellet was frozen at -20. Genomic DNA was extracted with the NucleoSpin Blood L kit. sgRNA cassettes were amplified, pooled, and sequenced as described [85]. CRISPR screens were analyzed using MAGeCK-iNC as previously described [85]. Briefly, raw sequencing reads were cropped and aligned using custom scripts that are publicly available (https://kampmannlab.ucsf.edu/resources). Raw phenotype scores and p-values were calculated for target genes and negative control genes using a Mann-Whitney U-test.

### Data Analysis

#### RNA-seq analysis

Sequencing data was aligned to the human reference genome hg38. Rbowtie2 was used to align and count the number of transcripts from aligned reads. Differentially expressed genes were determined using DEseq2.

#### ATAC-seq analysis

Sequencing data was aligned to the human reference genome hg38 using Rbowtie2. Peak calling was performed with MACS2. Differential ATACseq was performed using DEseq2, and motif analysis was performed with the motifDB and motifmatchr packages. Differential motif analysis was performed with the chromVar package.

#### Gene set enrichment analysis

Enrichr was used to perform gene set enrichment analysis on RNA-seq, proteomics and phosphoproteomics datasets [91]. Non-overlapping top significant terms were plotted. Overlapping significant terms are defined as two or more GO terms that are made up of identical gene sets. The term with the highest combined score from a set of overlapping terms was plotted to simplify the visualization and better represent the underlying data.

#### Proteomics and Phosphoproteomics Analysis

Raw files were searched using the directDIA+ feature in Spectronaut, with DDA files provided as supplementary search files against a full human proteome from Uniprot (reviewed entries only, isoforms included). Phosphosites were extracted from the PTMsites output table from Spectronaut, and collapsed using the Tukey’s median polish functionality of MSstats in R.

#### Longitudinal image analysis

Brightfield images were stitched first using FIJI’s Grid/Collection stitch plugin [92]. The configuration files were then formatted to be compatible with the MIST stitching plugin, which allows input configuration files to be applied to other images [93]. The LCK-mNeonGreen images were then stitched using the brightfield coordinates with MIST. Images were blinded with a simple python script written by a colleague that changed the names of the images to another number greater than what is found in a 96 well plate (ex: B02 becomes B68). Blinded images were then manually traced and measured in FIJI. The longest primary neurite was considered to be the axon. Branches from that axon were considered to be axon branches.

Additional neurites extending from the soma were considered secondary neurites. The measurement files were then unblinded by a complementary python script. The unblinded data was imported into R, processed, and then plotted in Prism for visualization. Statistical analysis was performed in Prism. One-way ANOVA with Dunnett’s multiple comparison test was performed on single timepoint data.

## Abbreviations

*MAPT*: Microtubule Associated Protein Tau
iPSC: induced pluripotent stem cell
MAPK: Mitogen-Activated Protein Kinase
MAP2K: Mitogen-Activated Protein Kinase Kinase
ASOs: Anti-sense oligonucleotides
FTD: Frontotemporal Dementia
WT: wild type
*MAPT* Het: *MAPT* heterozygous (*MAPT* V337M/WT)
*MAPT* Hom: *MAPT* homozygous (*MAPT* V337M/V337M)
mNGN2: mouse Neurogenin2
AAVS1: adeno-associated virus integration site 1, safe harbor locus.
CLYBL: Citramalyl-CoA Lyase, here refers to an intergenic safe harbor locus.
sgRNA: single guide RNA
NTC: non-targeting control
*ANK3*: Ankyrin 3
*MAPRE3*: Microtubule-Associated Protein RP/EB Family Member 3, or EB3
*GSK3B*: Glycogen Synthase Kinase 3 Beta
*CDK5*: Cyclin Dependent Kinase 5
*CDK5R1*: Cyclin Dependent Kinase 5 Regulatory Subunit 1
*MARK1*: Microtubule Affinity Regulating Kinase 1
*Map1a*: Microtubule-Associated Protein 1A
*Map1b*: Microtubule-Associated Protein 1B
LDEV: Lactate Dehydrogenase Elevating Virus
DMEM: Dulbecco’s Modified Eagle Medium
ROCK: Rho Associated Coiled-Coil Containing Protein Kinase
BDNF: Brain-Derived Neurotrophic Factor
KRAB: Krüppel-associated box
TALENS: transcription activator-like effector nucleases
PBS: Phosphate-buffered saline
TBST: Tris-buffered saline + 0.1% Tween20
TCEP: Tris 2-carboxyethyl phosphine
CAA: chloroacetamide
TFA: trifluoroacetic acid
ACN: acetonitrile
FA: formic acid
KFF: King Fisher Flex
DDA: Data dependent analysis
AGC: Automated gain control
DIA: Data-independent analysis
MS: Mass spectrometry
Ppm: parts per million
PTM: Post translational modification
MS/MS: Tandem mass spectrometry
HCD: Higher energy collisional dissociation
*GAPDH*: glyceraldehyde-3-phosphate dehydrogenase
*PSAP*: Prosaposin
emGFP: emerald green fluorescent protein
*BRD2*: Bromodomain Containing 2
ORF: open reading frame
TMP: trimethoprim
PDL: poly-D lysine
FACS: fluorescence-activated cell sorting
MAGeCK-iNC: Model-based Analysis of Genome-wide CRISPR/Cas9 Knockout – including negative controls

## Declarations

### Ethics approval and consent to participate

Studies at UCSF with human iPSCs were approved by the Human Gamete, Embryo, and Stem Cell Research (GESCR) Committee.

### Consent for publication

All authors have approved the contents of this study and provided consent for publication.

## Competing interests

M.K. is a co-scientific founder of Montara Therapeutics and serves on the Scientific Advisory Boards of Engine Biosciences, Casma Therapeutics, Cajal Neuroscience, Alector, and Montara Therapeutics, and is an advisor to Modulo Bio and Recursion Therapeutics. M.K. is an inventor on US Patent 11,254,933 related to CRISPRi and CRISPRa screening, and on a US Patent application on *in vivo* screening methods.

## Funding

M.K. was supported by the Rainwater Charitable Foundation/Tau consortium, the Chan Zuckerberg Initiative Ben Barres Early Career Acceleration Award, and NIH grants R01 AG062359, R01 AG082141, U54 NS100717. M.K. and D.L.S were supported by U54 NS123746. G.D. was supported by the A.P. Gianni Foundation. A.J.S was supported by NIH F32 AG063487 and NIH K99 AG080116-01. S.C.B. was supported by the Alzheimer’s Association Research Fellowship 23AARF-1027616.

## Author’s contributions

G.A.M., G.D. and M.K. conceptualized and led the project and wrote the manuscript with input from all co-authors. G.A.M, G.D., E.M., A.J.S., C.P.S, J.J., N.A., A.K., C. C., A.L., and S.C.B. contributed to making cell lines, performing experiments, and data analysis. J.M. and D.P.S. performed and analyzed the mass spectrometry.

## Acknowledgements

We would like to thank all members of the Kampmann lab for helpful feedback and support. We would like to thank Dr. Ruilin Tian, Dr. Carlo Condello, Dr. Noam Teyssier, Dr. Parker Grosjean, Dr. Olivia Teter and Ian Steele for helpful discussions and contributions to preliminary studies. We would like to thank the Tau Consortium Stem Cell Group for sharing the GIH6C1 and GIH6C1Δ1E11 iPSC cell lines and Dr. Li Gan for sharing the WTC11 V337M iPSC cell lines.

## Authors’ Information

Gregory A. Mohl and Gary Dixon have contributed equally to this work.

---

**Figure S1:**
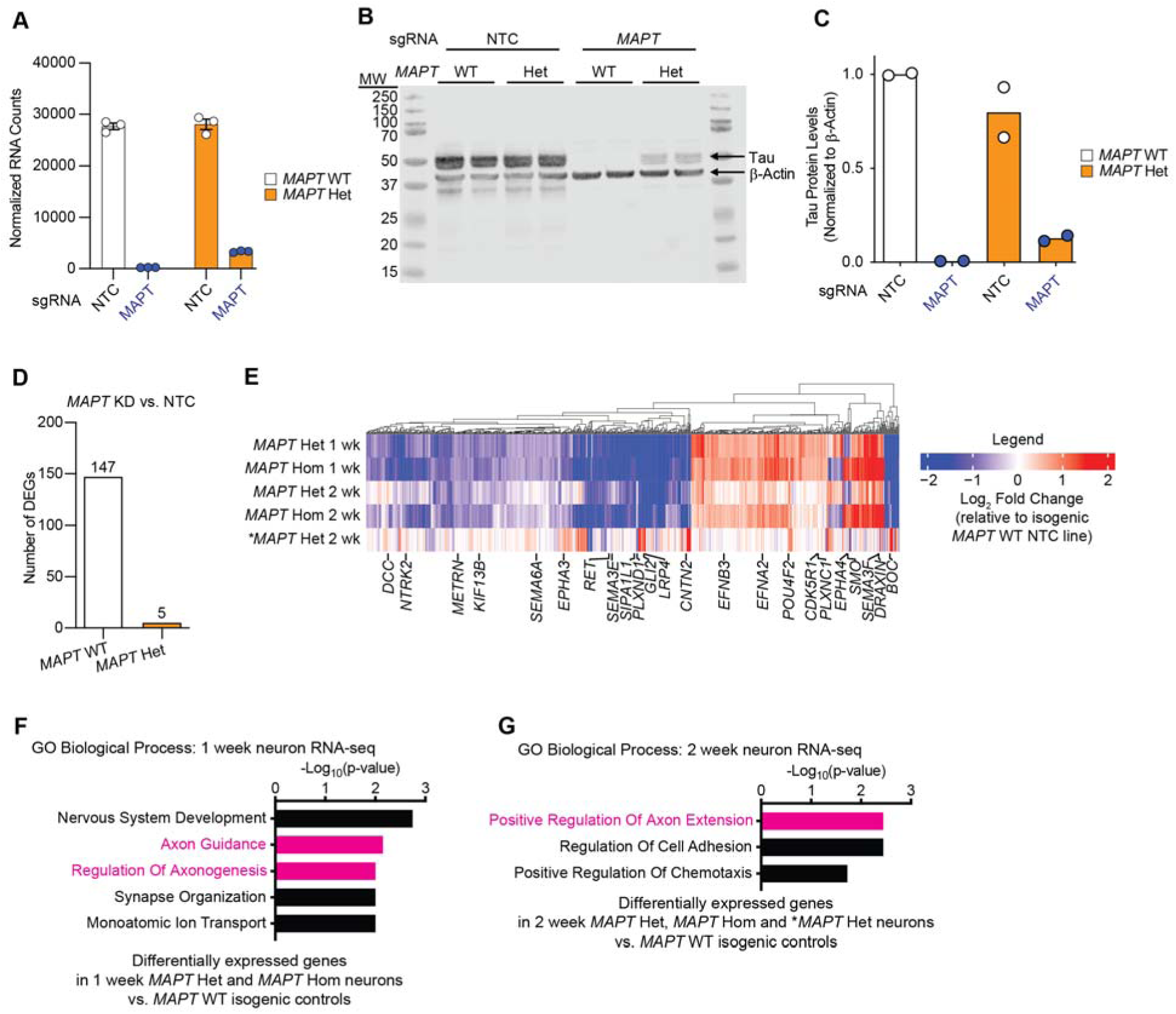
The *MAPT* V337M mutation and *MAPT* knockdown perturb gene expression of axonogenesis-related genes. **(A)** Normalized RNA counts of *MAPT* from the RNA-seq experiment described in Figure 1B showing tau knockdown in *MAPT* WT and *MAPT* Het neurons. **(B)** Western blot measuring tau knockdown in *MAPT* WT and *MAPT* Het neurons. Two replicates (individual wells) of neurons were harvested after two weeks of differentiation. **(C)** Quantification of the western blot in (B). **(D)** Bar plot showing the number of differentially expressed genes due to *MAPT* KD in either *MAPT* WT or *MAPT* Het neurons. **(E)** Heatmap of RNA-seq from *MAPT* Het, *MAPT* Hom and \**MAPT* Het neurons vs. isogenic controls at 1 week or 2 weeks of differentiation. Differentially expressed genes related to axon guidance or axonogenesis are labeled. **(F)** GO term enrichment analysis of one-week neurons from the RNA-seq experiment in (A). Genes that are differentially expressed in both *MAPT* Het and *MAPT* Hom vs. *MAPT* WT were analyzed with Enrichr, and top terms were plotted. Pathways related to axonogenesis and neuron morphology are colored magenta. (G) GO term enrichment analysis of two-week old neurons from the RNA-seq experiment in (A). Genes that are differentially expressed in both *MAPT* Het, *MAPT* Hom and \**MAPT* Het vs. their isogenic *MAPT* WT controls were analyzed with Enrichr, and top terms were plotted. Pathways related to axonogenesis and neuron morphology are colored magenta.

**Figure S2:**
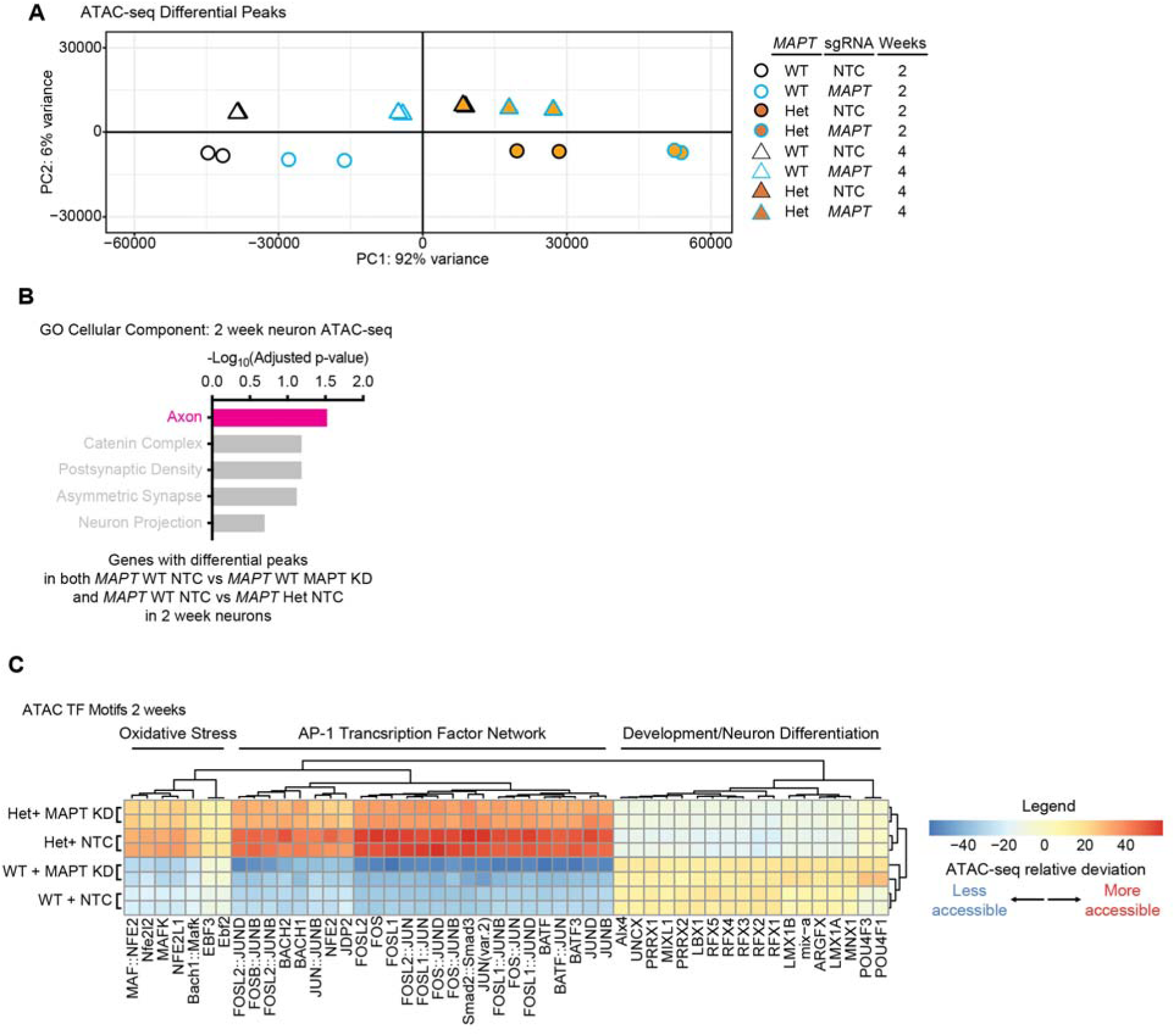
The *MAPT* V337M mutation and *MAPT* knockdown perturb chromatin accessibility of AP-1 transcription factor network motifs. **(A)** PCA plot of ATAC-seq differential peaks at 2 and 4 weeks of differentiation. Two replicates (individual wells) of neurons were harvested at each timepoint. **(B)** GO term enrichment analysis using Cellular Component on genes in 2-week neurons with differential ATAC-seq peaks in both *MAPT* WT *MAPT* KD and *MAPT* Het NTC vs. *MAPT* WT NTC. The top five terms were plotted, and non-significant terms are labeled in grey. **(C)** Heatmap showing the relative deviation of transcription factor motifs with significantly different accessibility in *MAPT* WT and *MAPT* Het neurons +/-tau knockdown. Two replicates (individual wells) of neurons were harvested at two weeks of differentiation. GO term enrichment analysis was used on clusters of transcription factors to categorize clusters. **(D)** Western blots for cJun and p-cJun in neurons at one week of differentiation. For each genotype, two independent differentiations with a total of five wells of neurons were analyzed for cJun, p-cJun and GAPDH levels.

**Figure S3:**
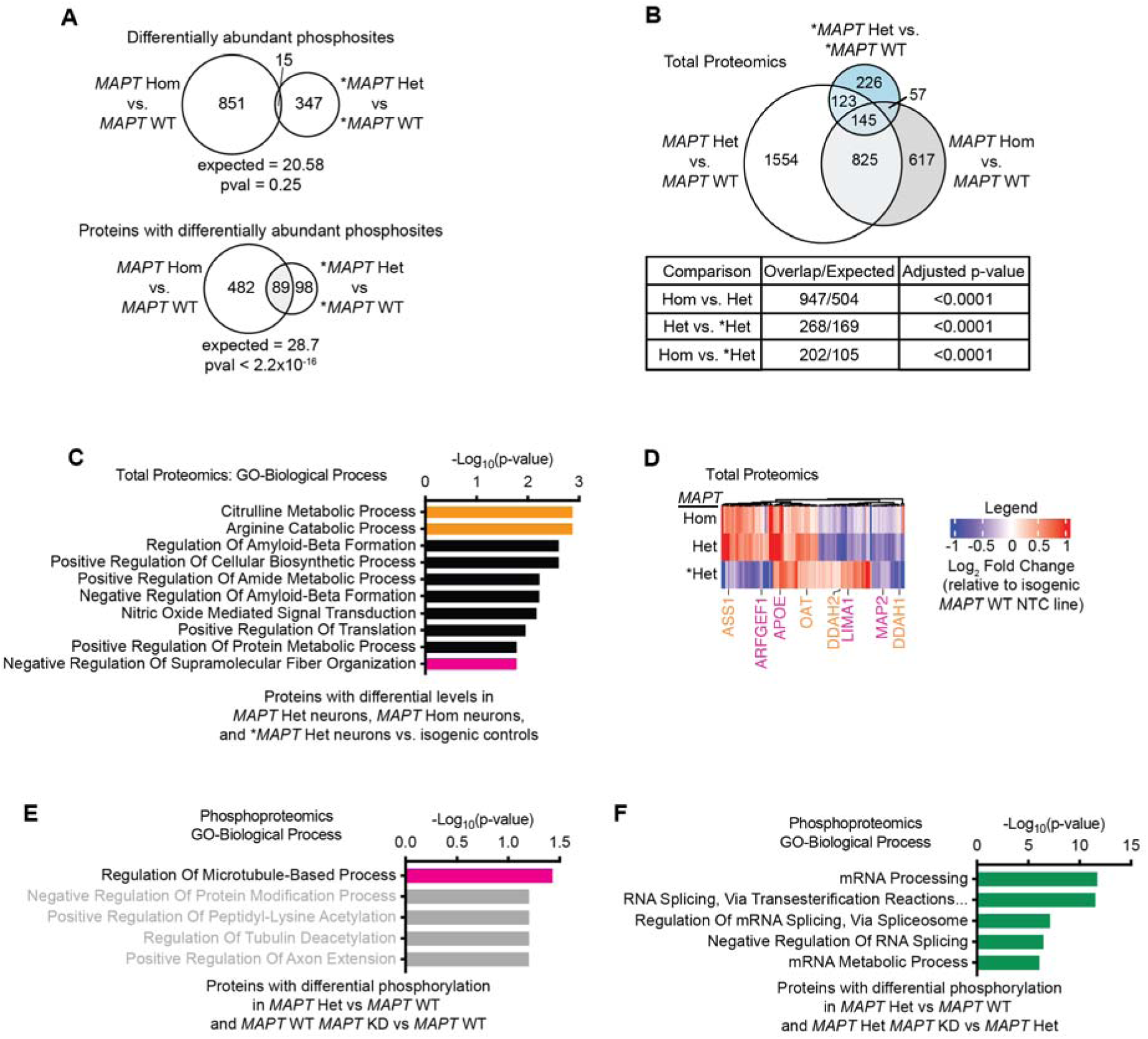
The *MAPT* V337M mutation and *MAPT* knockdown cause phosphorylation changes in axonogenesis and splicing proteins. **(A)** (*Top*) Overlap between differential phosphosites in neurons derived from iPSCs edited to introduce the homozygous *MAPT* V337M mutation (*MAPT* Hom)vs. isogenic controls (*MAPT* WT). Four replicates (independent 150mm dishes) of neurons for each genotype/sgRNA combination were harvested after one week of differentiation, and the phosphoproteome was measured using mass spectrometry. Significance was calculated using Fisher’s Exact Test. (*Bottom*) Overlap between proteins with differential phosphorylation in both datasets. Significance was calculated using Fisher’s Exact Test. **(B)** Overlap between proteomic changes in *MAPT* Hom neurons, neurons derived from iPSCs edited to have the heterozygous *MAPT* V337M mutation (*MAPT* Het) and neurons derived from patient iPSCs with the heterozygous *MAPT* V337M mutation (*\*MAPT* Het) vs. isogenic controls (*MAPT* WT or \**MAPT* WT). Four replicates (independent 150mm dishes) of neurons for each genotype/sgRNA combination were harvested after one week of differentiation, and the total proteome was measured using mass spectrometry. Significance was calculated using multiple t-tests adjusted with Šidák single-step correction. Significantly differential proteins in all three datasets were filtered to identify 145 conserved proteins. **(C)** GO term enrichment of the 145 proteins with differential abundance in *MAPT* Hom, *MAPT* Het and \**MAPT* Het neurons compared to isogenic controls. Top non-overlapping significant terms are shown. Term names are colored to match relevant gene names in the heatmap in (C). **(D)** Heatmap showing the Log_2_ fold change of protein abundance for the 145 proteins with differential abundance in *MAPT* Hom, *MAPT* Het and \**MAPT* Het neurons vs. isogenic *MAPT* WT neurons. Proteins within enriched GO terms are labeled and colored according to the shared pathways. **(E)** GO term analysis of phospoproteins with differential phosphorylation in *MAPT* Het NTC and *MAPT* WT *MAPT* KD vs. *MAPT* WT NTC. Non-significant terms are labeled by grey bars. Regulation of Microtubule-based process is labeled by a magenta bar due to its overlap with axon-related terms. **(F)** GO term analysis of phosphoproteins with differential phosphorylation in *MAPT* Het NTC vs. *MAPT* WT NTC and *MAPT* Het *MAPT* KD vs. *MAPT* Het NTC. Terms related to RNA processing and splicing are marked by green bars.

**Figure S4:**
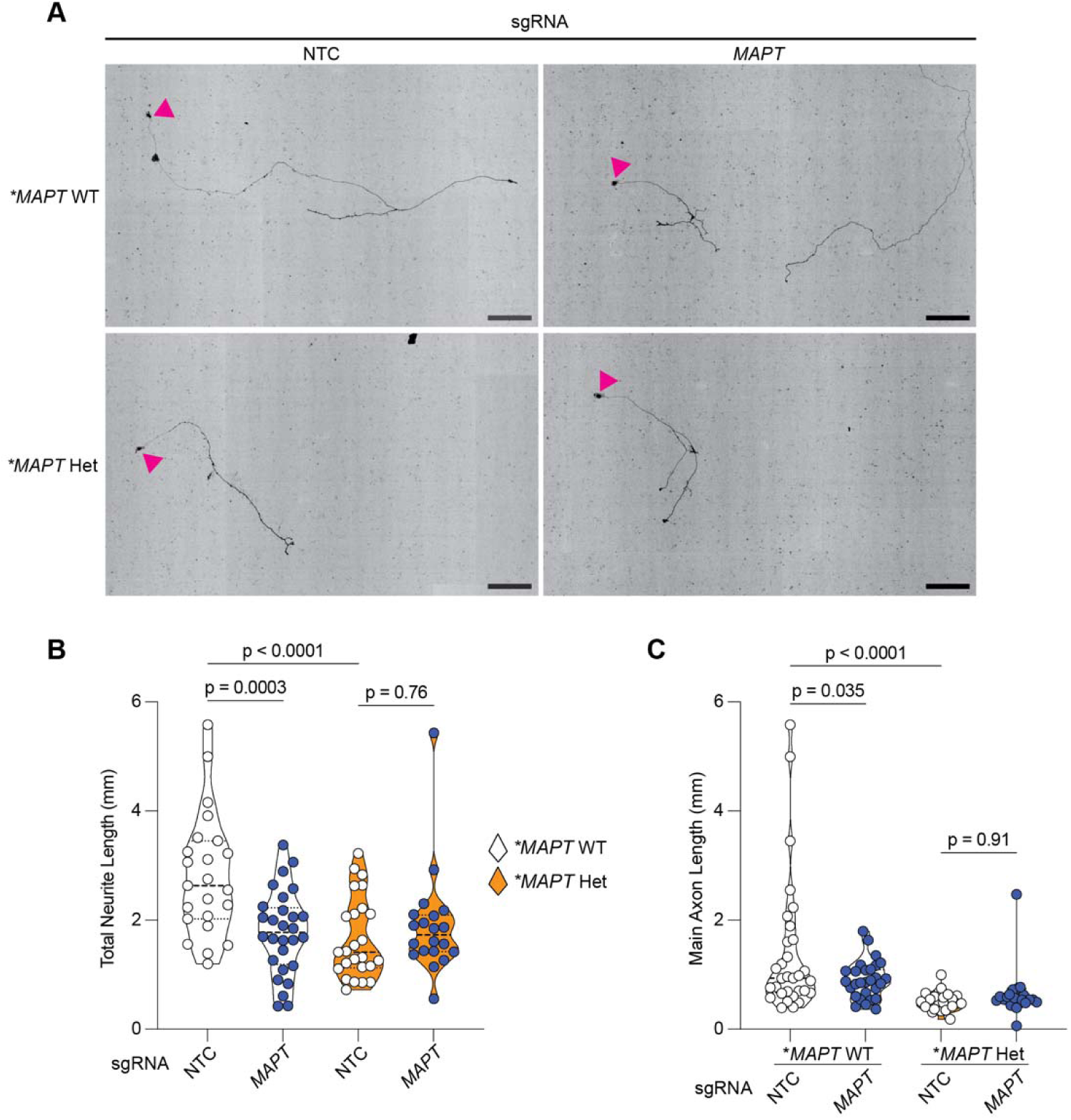
The *MAPT* V337M mutation and *MAPT* knockdown decrease main axon length and total neurite length in patient-derived neurons. **(A)** Representative images of neurons labeled with mNeonGreen. The scale bars are 200µm. Magenta arrowhead indicates the nucleus. **(B-C)** Quantification of total neurite length (B) and main axon length (C). Significance was calculated using one-way ANOVA with Šidák’s multiple comparison test.

**Figure S5:**
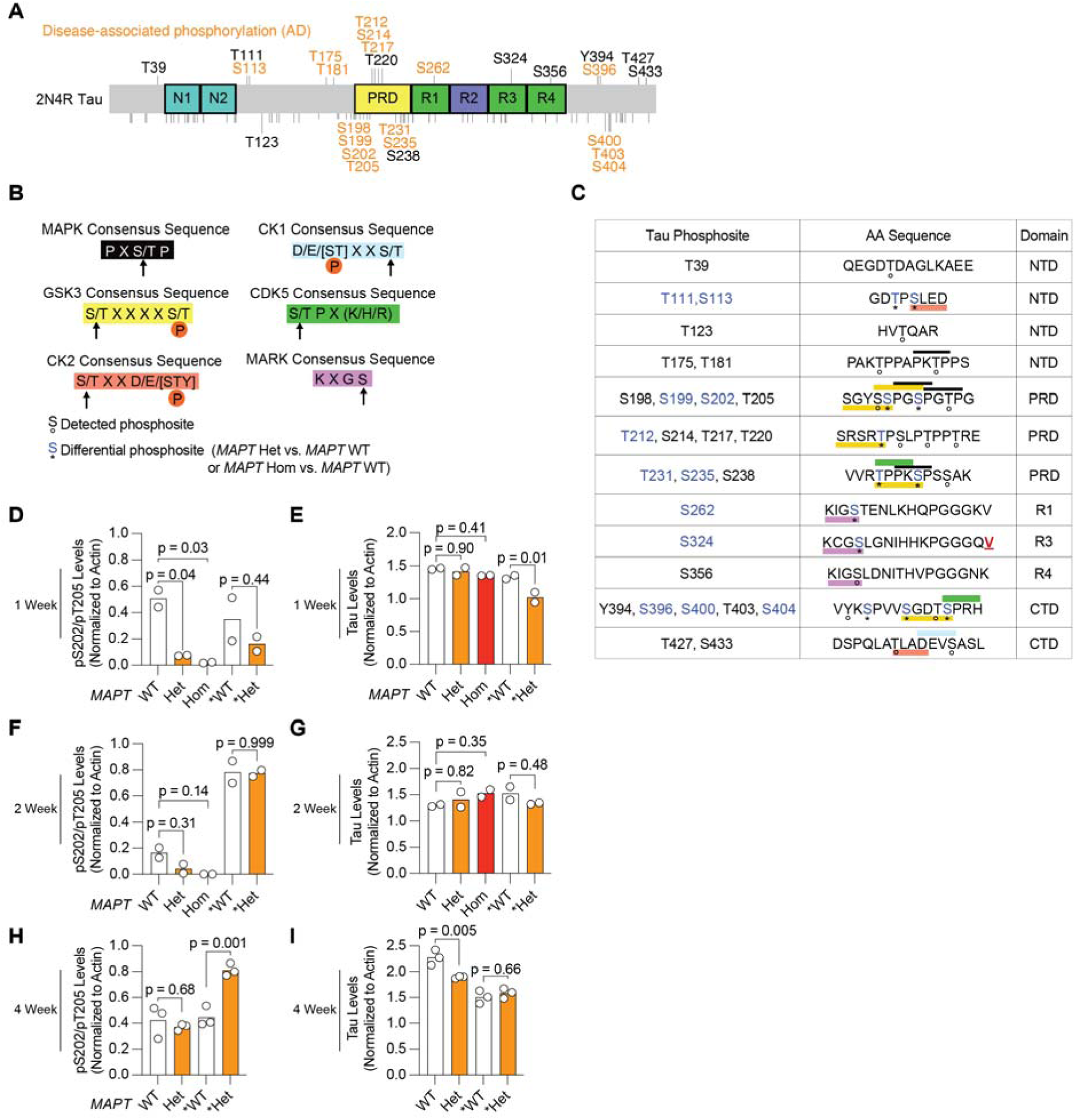
Neurons with the *MAPT* V337M mutation have decreased tau phosphorylation at disease-associated phosphorylation sites. **(A)** Protein domain map of 2N4R tau. Phosphosites detected in this study are labeled, with disease-associated phosphorylation sites from AD labeled in orange. Phosphosites not detected in this study are marked with a small black line and are unlabeled. Domain abbreviations are as follows: N-terminal inserts (N1,N2), proline rich domain (PRD), microtubule binding repeats (R1, R2, R3, R4). **(B)** Consensus sequences for tau kinases. Kinase consensus sequences are annotated with colored boxes, with priming sites marked with a “P” in an orange circle. **(C)** Detected tau phosphosites are shown with their sequence context. Phosphorylation sites that are differential between either *MAPT* V337M heterozygous (*MAPT* Het) or *MAPT* V337M homozygous (*MAPT* Hom) are labeled blue with an asterisk, and detected phosphosites are labeled with an open circle. V337 is labeled with a bold/underlined red V. The domains abbreviated as follows: N-terminal projection domain (NTD), proline rich domain (PRD), Microtubule binding repeats (R1, R3, R4), C-terminal domain (CTD). **(D-I)** Quantification of western blots in Figure 3C. Significance was calculated using one-way ANOVA with Šidák’s multiple comparison test.

**Figure S6:**
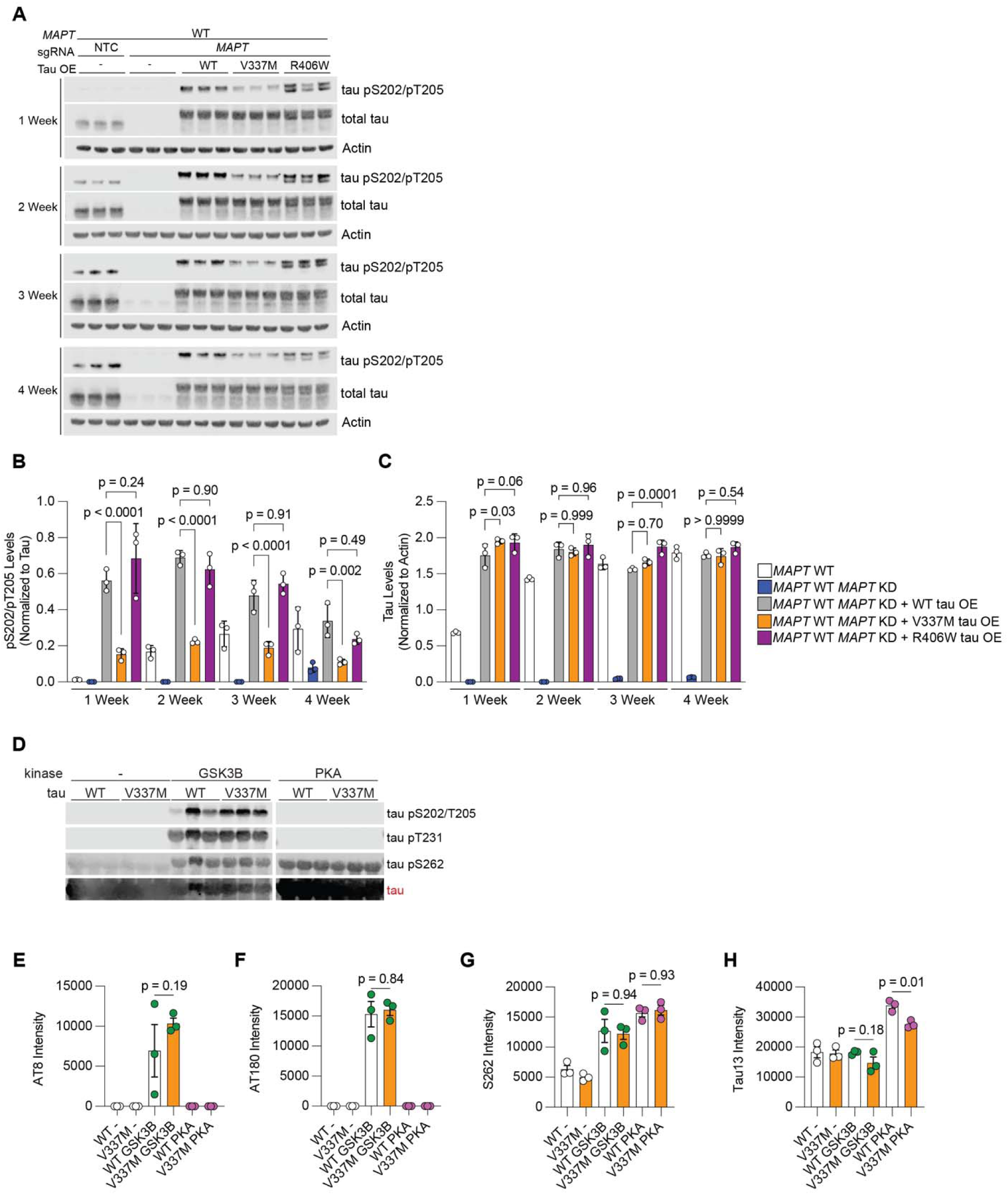
V337M tau is hypophosphorylated when overexpressed in *MAPT* WT *MAPT* KD neurons. **(A)** Timecourse of tau pS202/pT205 and tau levels in neurons overexpressing WT, V337M or R406W tau. **(B-C)** Quantification of tau pS202/pT205 levels (B) and total tau levels (C) from the blots in (A). Significance was calculated using one-way ANOVA with Šidák’s multiple comparison test. **(D)** Western blot measuring *in vitro* tau phosphorylation with GSK3B or PKA. (E-G) Quantification of western blots in (D). Significance was calculated using one-way ANOVA with Šidák’s multiple comparison test.

**Figure S7:**
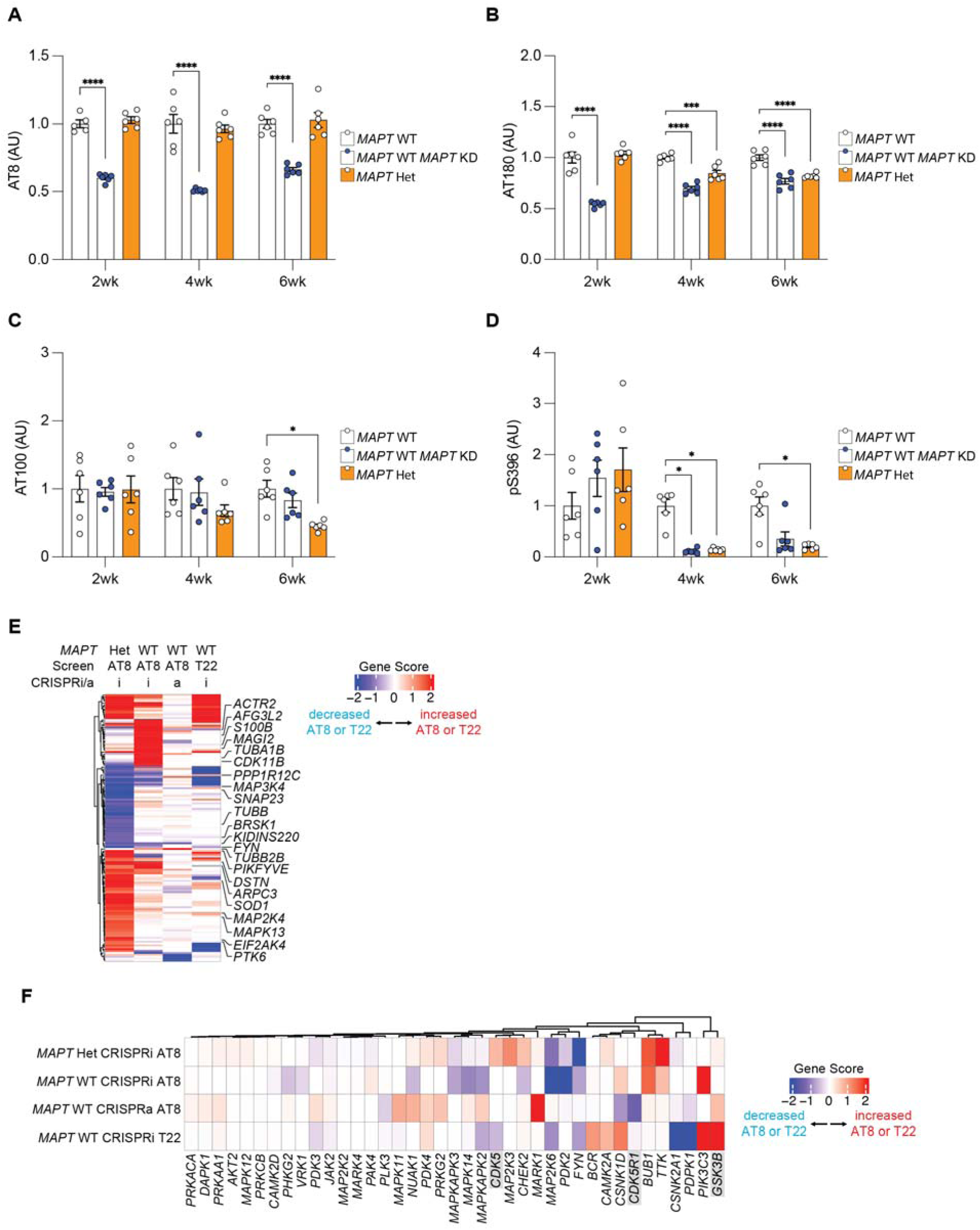
Functional genomics uncovers regulators of tau phosphorylation in *MAPT* WT and *MAPT* Het neurons. (A-D) Bar plots showing the median intensity of AT8 (A), AT180 (B), AT100 (C) and pS396 (D) in 2-week *MAPT* WT, *MAPT* KD and *MAPT* Het neurons. AT8 was selected for CRISPR screening due to high reproducibility across timepoints and AT8 detection in both *MAPT* WT and *MAPT* Het neurons at 2 weeks of differentiaton. **(E)** Heatmap of hits from the CRISPRi and CRISPRa AT8 screens and the CRISPRi T22 screen. Many of the AT8 hits from the three screens do not modify T22 levels and are therefore unlikely to be due to modifying tau levels [90]. Genes related to cytoskeleton, neuron projection development or the p38 MAPK pathway are annotated. **(F)** Heatmap of AT8 and T22 screens with the kinases predicted to have differential activity in Figure 3I. Selected kinases predicted to have differential activity in *MAPT* V337M neurons with particular disease relevance that did not have a phenotype in the AT8 screens are highlighted with grey boxes.

**Figure S8:**
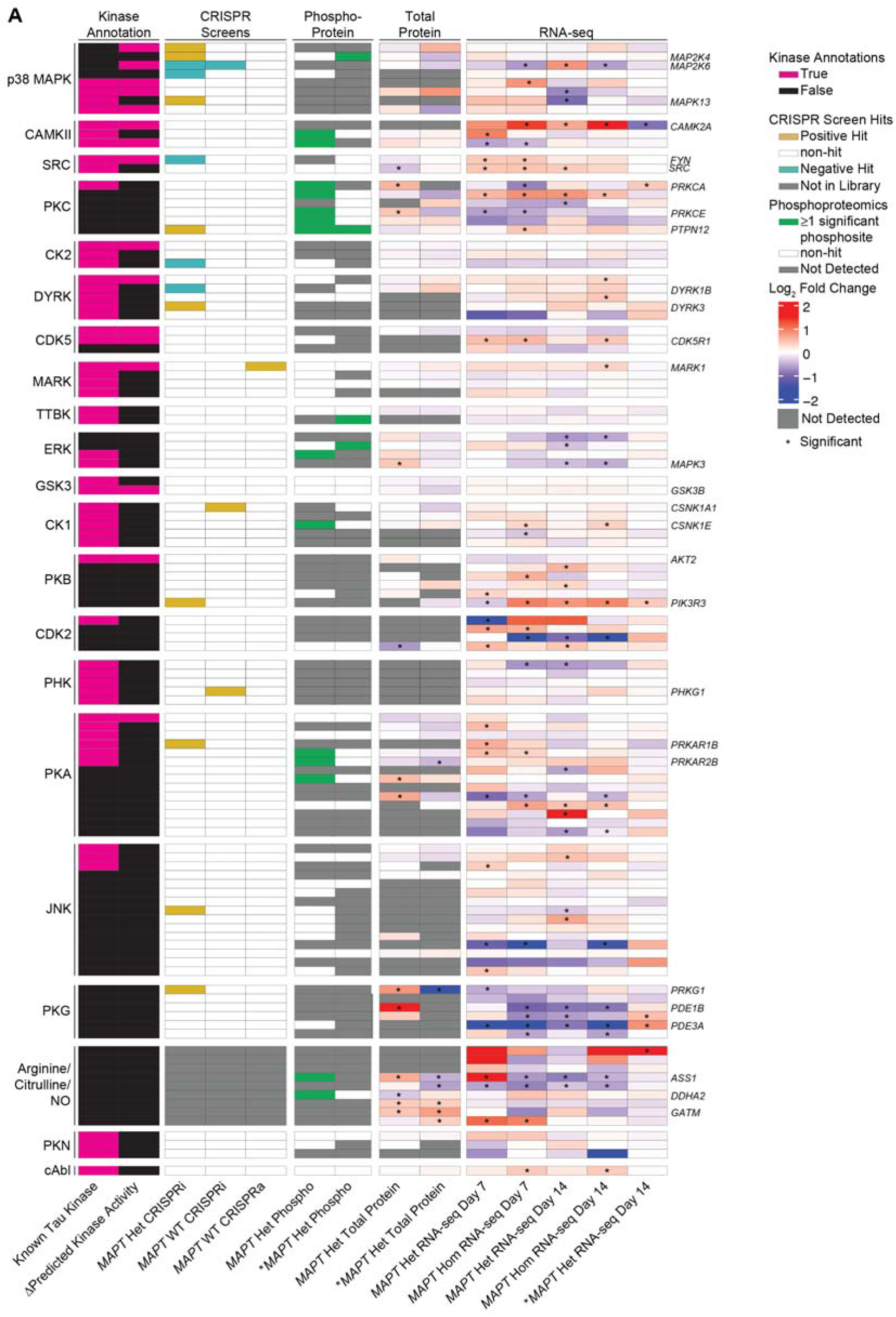
The p38 MAPK pathway regulates tau phosphorylation in *MAPT* Het neurons. **(A)** Heatmap showing known tau kinases compared to the predicted kinase activity in *MAPT* V337M neurons, CRISPRi/a screens, phosphoproteomics, total proteomics and transcriptomics. Several kinases in the p38 MAPK pathway modulate pS202/pT205 phosphorylation, though most were not detected in the phosphoproteomics. Most of these genes were not differentially expressed at the protein or RNA level. PKG and the Arginine/Citrulline genes were included because many of these genes were differentially expressed at the RNA and protein level as a counter example to the p38 MAPK pathway.

**Figure S9:**
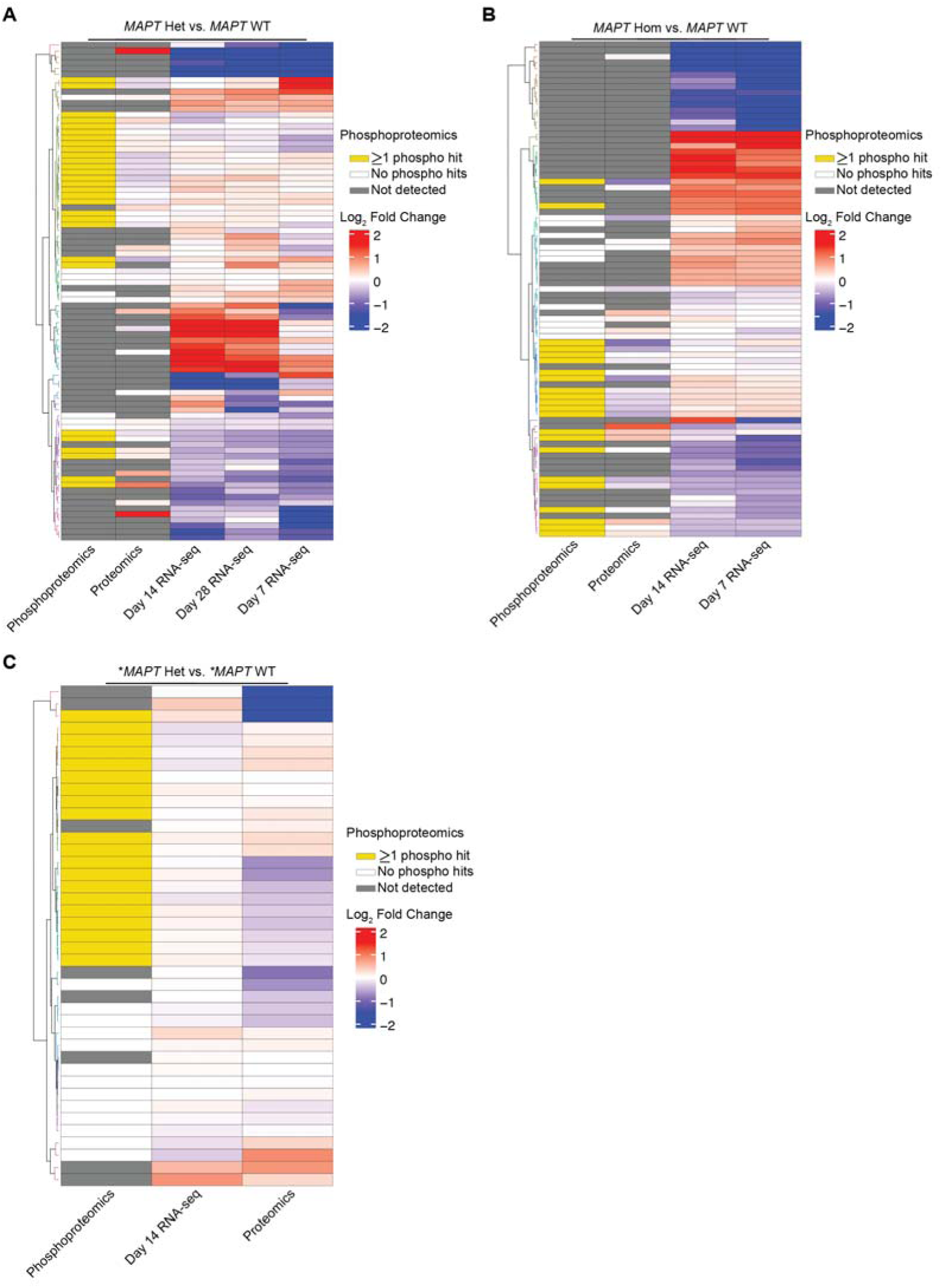
Most differentially phosphorylated proteins involved in axonogenesis in V337M neurons are not differentially expressed at the mRNA or total protein levels. **(A)** Heatmap showing the phosphoproteomics, total proteomics, and RNA-seq data for genes involved in axonogenesis pathways in *MAPT* Het vs. *MAPT* WT neurons. **(B)** Heatmap showing the phosphoproteomics, total proteomics, and RNA-seq data for genes involved in axonogenesis pathways in *MAPT* Hom vs. *MAPT* WT neurons. **(C)** Heatmap showing the phosphoproteomics, total proteomics, and RNA-seq data for genes involved in axonogenesis pathways in \**MAPT* Het vs. \**MAPT* WT neurons.

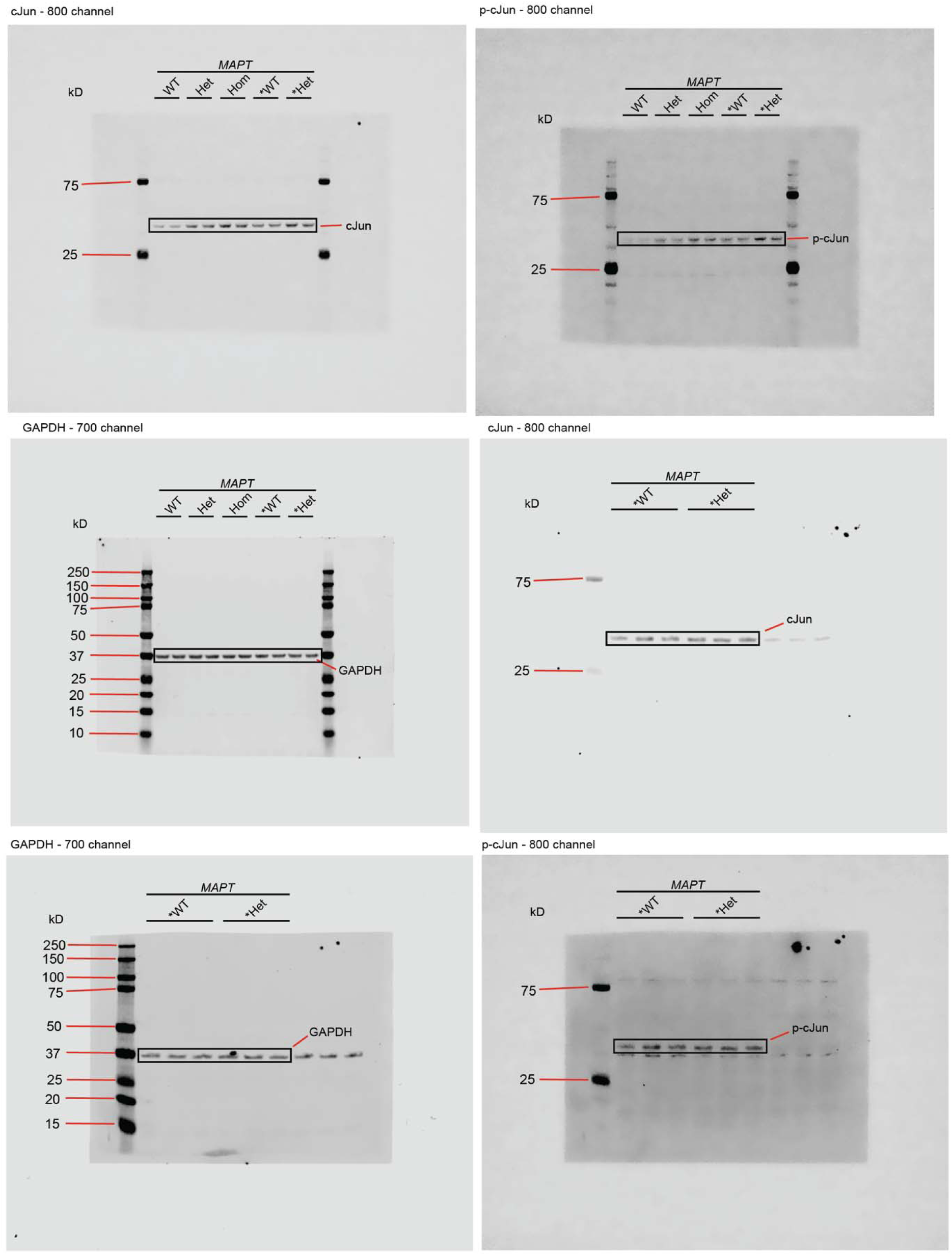
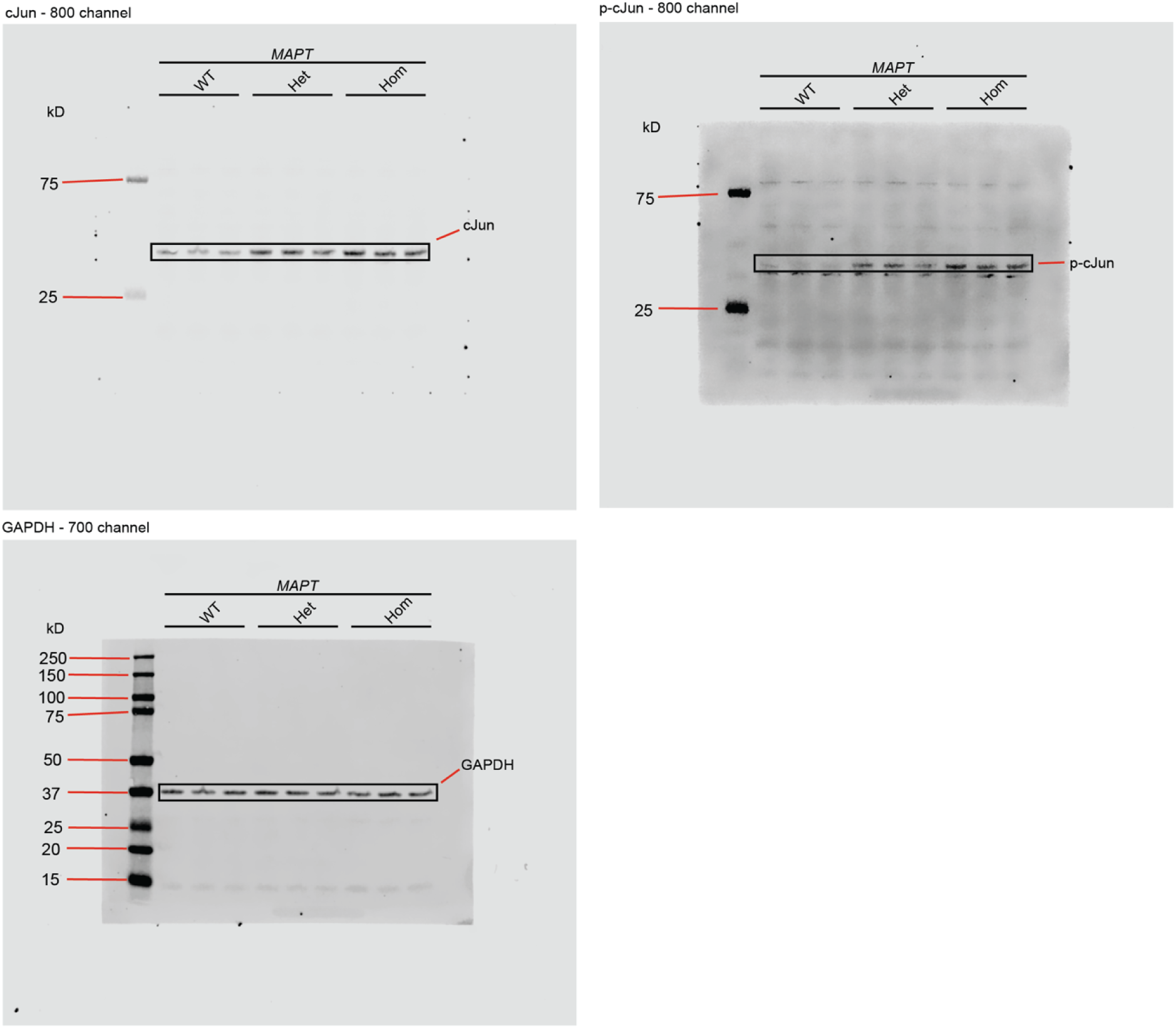

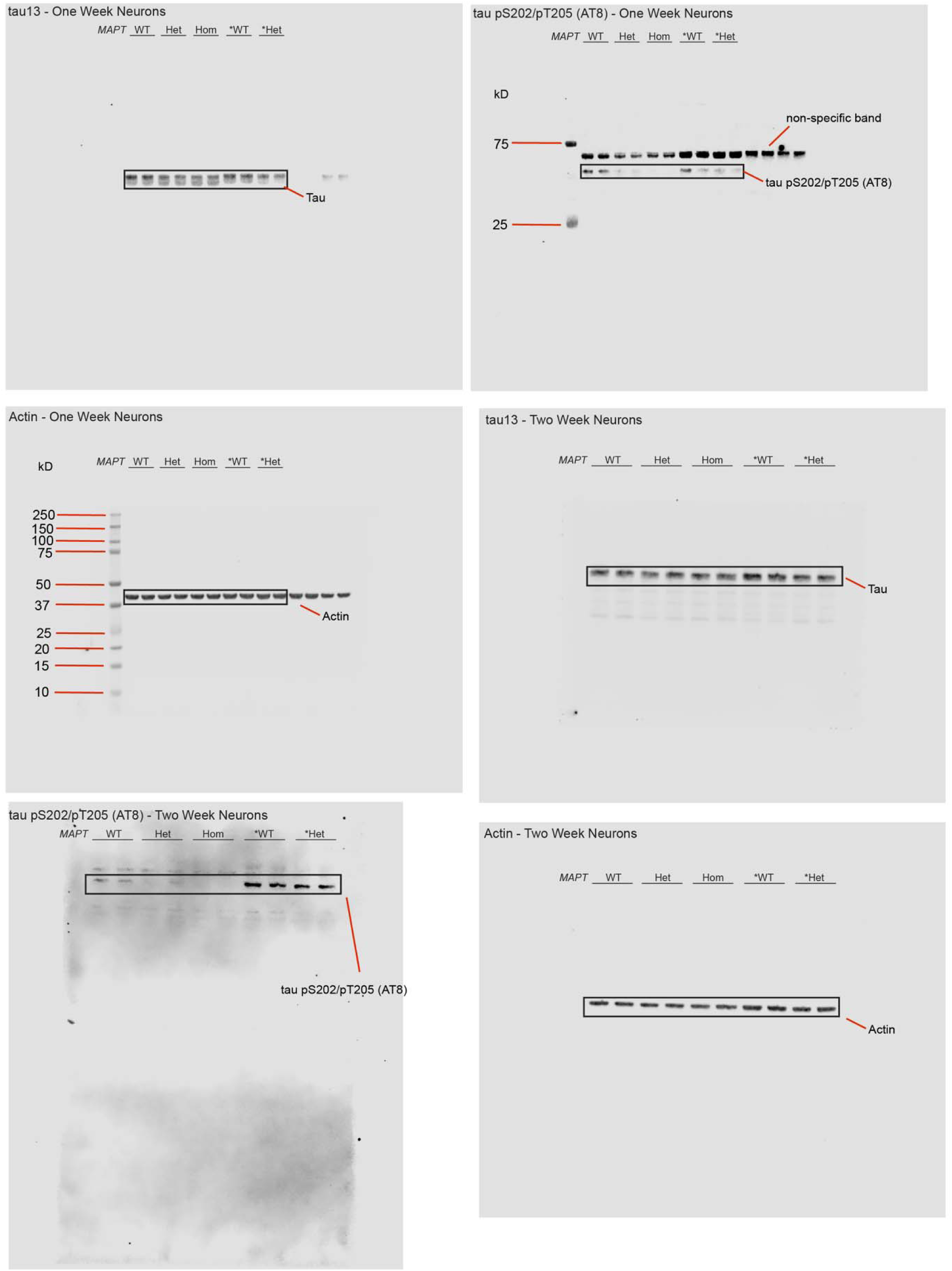
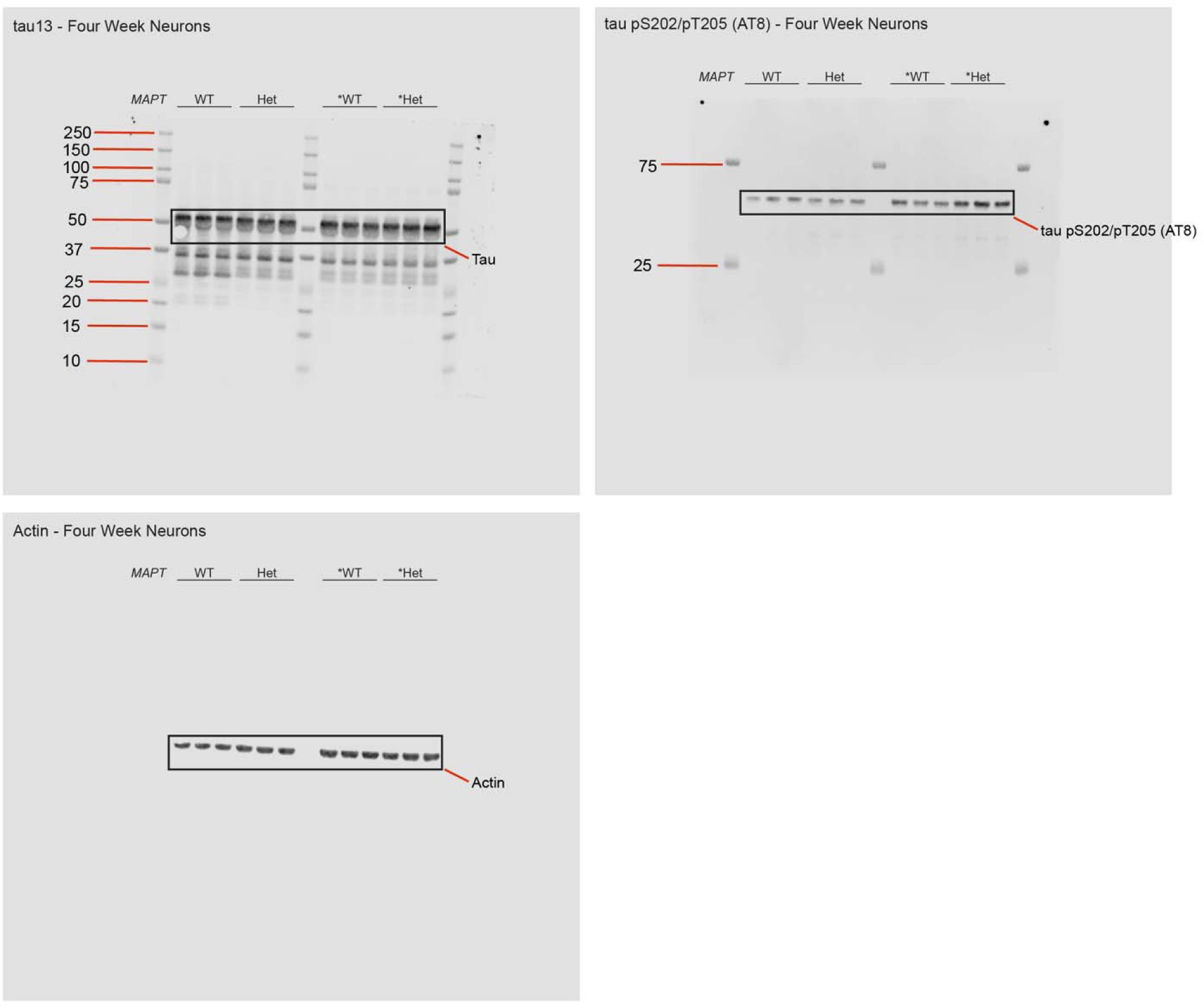

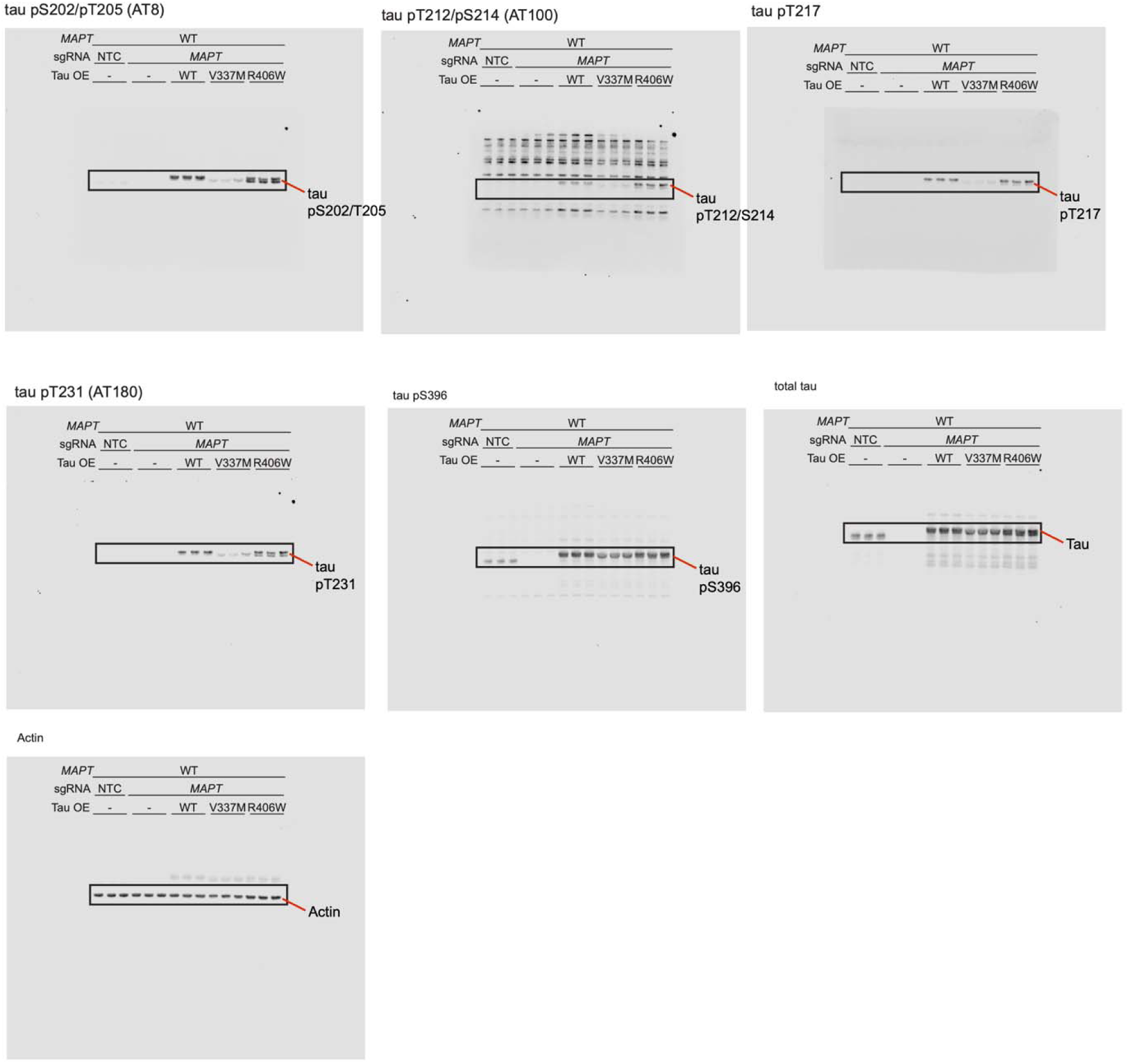

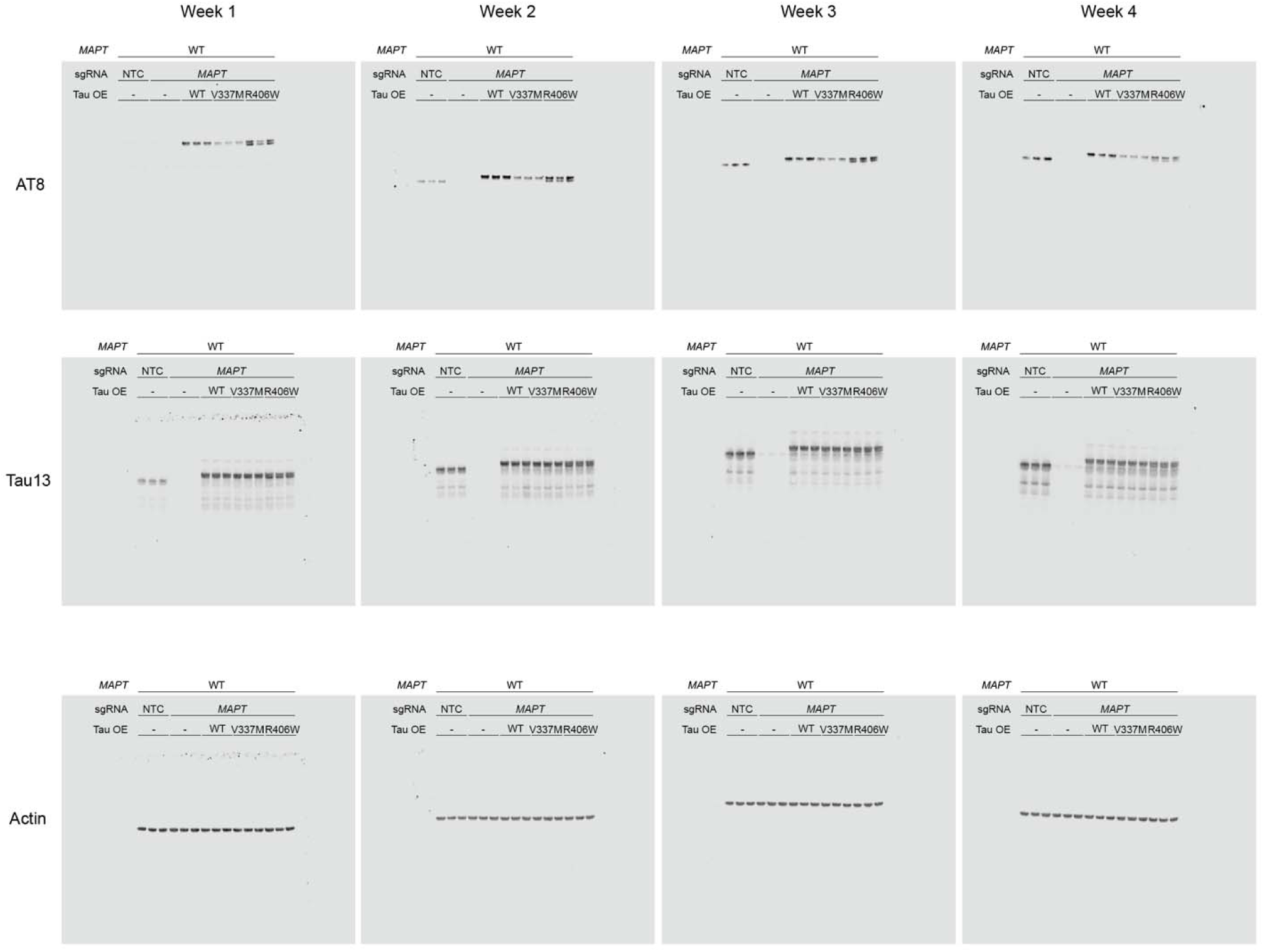

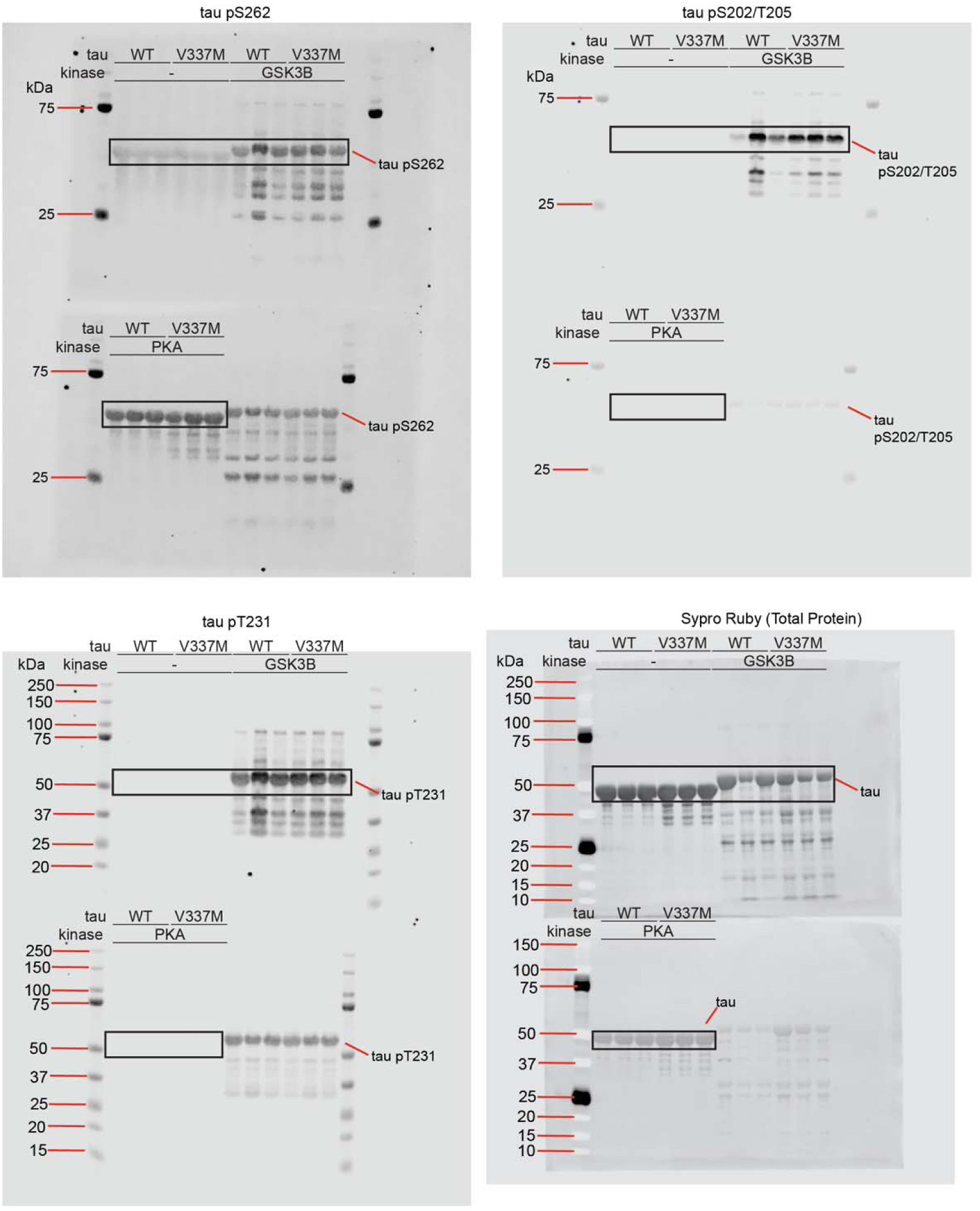

## Notes

### Summary of Updates

Additional experiments, analyses and discussion. 2 additional authors.

